# Structure of the human outer kinetochore KMN network complex

**DOI:** 10.1101/2023.08.07.552234

**Authors:** Stanislau Yatskevich, Jing Yang, Dom Bellini, Ziguo Zhang, David Barford

## Abstract

Faithful chromosome segregation requires robust, load-bearing attachments of chromosomes to the mitotic spindle, a function accomplished by large macromolecular complexes termed kinetochores. In most eukaryotes, the constitutive centromere-associated network (CCAN) complex of the inner kinetochore recruits to centromeres the ten-subunit outer kinetochore KMN network, which comprises the KNL1C, MIS12C and NDC80C complexes. The KMN network directly attaches CCAN to microtubules through MIS12C and NDC80C. Here, we determined a high-resolution cryo-EM structure of the human KMN network. This showed an intricate and extensive assembly of KMN subunits, with the central MIS12C forming rigid interfaces with NDC80C and KNL1C. The redundancy and strength of inter-subunit connections explains how KMN withstands strong forces applied during chromosome segregation. We also observed that unphosphorylated MIS12C exists in an auto-inhibited state that suppresses its capacity to interact with CCAN. Ser100 and Ser109 of the N-terminal segment of the MIS12C subunit Dsn1, two key targets of Aurora B kinase, directly stabilize this auto-inhibition. Our work provides a molecular mechanism for how selectively relieving this auto-inhibition through Ser100 and Ser109 phosphorylation would restrict outer kinetochore assembly to functional centromeres during cell division.

## Introduction

Kinetochores are large macromolecular complexes that couple the forces of microtubule depolymerization to chromosome movement during mitosis and meiosis^1–3^. The chromatin-proximal inner kinetochore, also known as the constitutive centromere associated network (CCAN), assembles specifically at the centromere by recognizing the centromere-specific histone H3 variant, CENP-A. Inner kinetochores together with CENP-A nucleosomes (CENP-A^Nuc^) form load-bearing attachments at centromeric chromatin and recruit the outer kinetochore which directly couples CCAN to microtubules of the mitotic spindle. While CCAN is present at the centromere throughout the cell cycle, in metazoans the outer kinetochore is recruited to centromeres only during mitosis^4^. Interactions between CCAN components and outer kinetochore components have to be sufficiently robust to withstand the pulling forces applied by depolymerizing microtubules, estimated to be approximately 1000 pN per chromosome^5^. The outer kinetochore also scaffolds the spindle assembly checkpoint (SAC), a critical signalling pathway that ensures accurate chromosome segregation^6–8^.

While the inner kinetochore diverged throughout evolution, the outer kinetochore is compositionally and structurally conserved in most species as the primary force-coupling device between chromosomes and the mitotic spindle^9^. The outer kinetochore is made up of the central and evolutionary conserved 10-subunit KMN network complex comprising the MIS12 (Mis12, Dsn1, Nsl1, Pmf1), NDC80 (Ndc80, Nuf2, Spc24, Spc25) and KNL1 (Knl1, ZWINT) complexes as well as auxiliary microtubule-associated components, such as the SKA complex, that vary in composition throughout evolution. The rod-shaped MIS12C functions as a central interaction hub by directly binding the Spc24:Spc25 RWD domains (Spc24:25^RWD^) of NDC80C as well as the Knl1 RWD domains (Knl1^RWD^) of KNL1C^10–13^, thereby coordinating the microtubule-binding (NDC80C) and checkpoint signalling modules (KNL1C) of the KMN network. MIS12C also constitutes one of the primary KMN network linkages to the centromere. At the bottom of the MIS12C stalk, two head domains interact with linear peptidic motifs of the inner kinetochore components – CENP-C and CENP-T^10, 11, 14–20^. Crystallographic studies of human and yeast CENP-C interactions with MIS12C showed that the N-terminal 45 residues of CENP-C contacts the head domain formed of the Mis12 and Pmf1 subunits (MIS12^Head-1^), while the head domain of Dsn1 and Nsl1 (MIS12^Head-2^) projects outwards^10, 11^. CENP-T binds MIS12C in a manner that competes with CENP-C binding and requires CENP-T phosphorylation by CDK1, suggesting that CENP-C and CENP-T bind a similar region of MIS12, but the molecular details of this interaction are less well understood^18, 19^.

NDC80C is the primary and essential force-coupling component of the kinetochore in nearly all eukaryotes^21^. NDC80C contains an approximately 50 nm long coiled-coil which separates the microtubule-binding calponin homology (CH) domains of Ndc80:Nuf2 at one end from Spc24:Spc25 interactions with MIS12C at the other^22–25^. Multimerization of NDC80C allows it to bind and track both growing and depolymerizing microtubules^21, 26^, a property that is dependent on a conserved coiled-coil loop (Ndc80^Loop^) that introduces a break into a continuous Ndc80:Nuf2 coiled-coil^27, 28^. The CCAN component CENP-T also recruits two copies of NDC80C in human cells, independently of the MIS12C pathway, by forming peptidic interactions with Spc24:25^RWD^ domains^18, 29, 30^.

Human Knl1 is a 2,342-residue protein predicted to be mostly unstructured. The only folded region of Knl1 is located at the C-terminus of the protein and contains Knl1^RWD^ and a predicted coiled-coil element, with Knl1^RWD^ being sufficient for Knl1 localization to kinetochores^13, 31^. The coiled-coil region of Knl1 is necessary to recruit its constitutive binding partner ZWINT^13^. The major function of Knl1 is to orchestrate SAC signalling via the recruitment of key SAC components such as Bub1, BubR1, and Bub3 using numerous conserved motifs^6^. ZWINT is also responsible for recruitment of the ROD:Zwilch:ZW10 (RZZ) complex that forms the fibrous corona around unattached kinetochores and also facilitates SAC signalling^32^. Additionally, the extreme N-terminus of Knl1 was shown to have an affinity for microtubules, suggesting that Knl1 also contributes to the formation of kinetochore-microtubule attachments^33, 34^.

Negative stain electron microscopy (EM) and rotary shadowing EM elucidated the global architecture of the KMN components as an elongated rod-like complex^11, 18^, yet the molecular details and the functional significance of how the constituent KMN complexes interact have remained elusive. Additionally, multiple studies suggest that MIS12C requires activation by Aurora B phosphorylation. Inhibiting Aurora B kinase activity disassembled kinetochores in Xenopus extracts^35^. In human cells, mutation of the two key Aurora B target residues, Ser100 and Ser109, located within an N-terminal extension of Dsn1 (Dsn1^N^), significantly impairs Dsn1 localization to kinetochores and compromises assembly of the functional outer kinetochore^36, 37^. Alanine substitution of the equivalent Ser100 and Ser109 residues in yeast is lethal and affects MIS12C recruitment to inner kinetochores, highlighting the functional importance and evolutionary conservation of these residues^38^. Biochemical studies suggested that Dsn1^N^ auto-inhibits the interaction of MIS12C with both CENP-C^10, 11, 36^ and CENP-T^19^, and that Aurora B kinase phosphorylation relieves this inhibition. However, the structural basis for Dsn1^N^-mediated auto-inhibition, and how this is relieved by phosphorylation to activate MIS12C binding to the inner kinetochore, is not understood.

To address these questions, we reconstituted the human KMN network and determined its structure using cryo-EM. We observed a rigid prong-shaped structure formed by the NDC80C:MIS12C:KNL1C components. Their mutual interactions are mediated by moderately-sized protein-protein interfaces that are further stabilized by extensive peptidic linkages, generating a configuration where NDC80C and KNL1C extend perfectly parallel with respect to one another from the central MIS12C scaffold. We also observed that MIS12C exists in an auto-inhibited state with MIS12^Head-1^ and MIS12^Head-2^ closely juxtaposed to one another. Dsn1^N^ directly stabilizes the interaction of the two MIS12C head domains, with Ser100 and Ser109 mediating auto-inhibiting interactions. In this binding mode, Dsn1^N^ competes with CENP-C and CENP-T for the CCAN-binding interface of MIS12, explaining how recruitment of KMN to the inner kinetochore might be regulated. We demonstrate that Aurora B-mediated phosphorylation would release the Dsn1^N^-mediated auto-inhibition, thereby regulating MIS12C:CENP-C interactions.

## Overall architecture of the KMN network junction

We purified the three sub-complexes of the KMN network, with all proteins apart from Knl1 being full-length (Extended Data Fig. 1a). For this study, we used an extended version of the Knl1 C-terminus encoding residues 1870-2342, which is sufficient for kinetochore localization and ZWINT binding^13^. Our Knl1 construct also contains all regions of Knl1 predicted to be folded by AlphaFold2^39^ (Extended Data Fig. 1b). We could reconstitute the complete KMN network using size exclusion chromatography (SEC), with all components co-eluting (Extended Data Fig. 1c). We subjected the reconstituted KMN network to cryo-EM analysis and obtained a 3.0 Å resolution reconstruction of the KMN network junction (KMN^Junction^) that contains the complete MIS12C, the complete Spc24:Spc25 proteins, and all folded domains of KNL1C (Fig. 1, Extended Data Fig. 2a-e, Extended Data Table 1). The only components we do not observe at high-resolution are the Ndc80 and Nuf2 proteins which appear to be flexibly connected to the Spc24:Spc25 proteins at the tetramerization junction.

**Figure 1.**
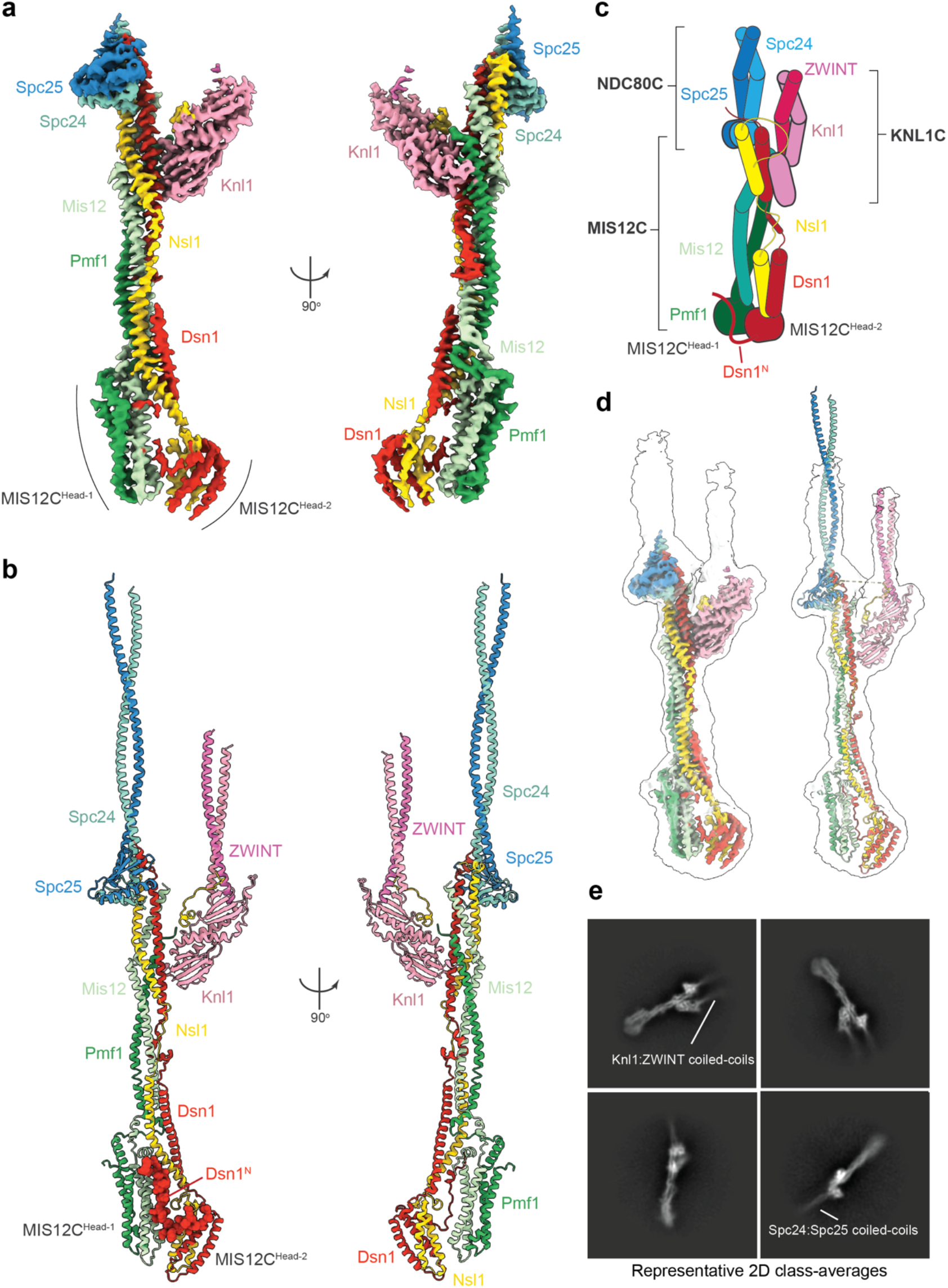
Overall architecture of the human KMN^Junction^ complex. a. Composite cryo-EM density map of the human KMN^Junction^ complex comprised of the rigid Spc24:Spc25:MIS12C:Knl1:ZWINT body and the more mobile MIS12^Head-1^:MIS12^Head-2^ body. b. Molecular model of the human KMN^Junction^ complex shown as a cartoon, highlighting the auto-inhibitory Dsn1^N^ region in space-filling representation. c. Cartoon schematic of the human KMN^Junction^ complex with diagrams of the multiple peptidic linkages present in the complex. d. A high-resolution sharpened composite map (left) and molecular model (right) of the human KMN^Junction^ complex fitted into the transparent grey consensus unmasked map of the KMN network complex that shows coiled-coil densities of the Spc24:Spc25 and Knl1:ZWINT complexes. e. Representative cryo-EM 2D class averages of the KMN network showing coiled-coils of Spc24:Spc25 and Knl1:ZWINT complexes projecting from the central KMN^Junction^ complex.

**Figure 2.**
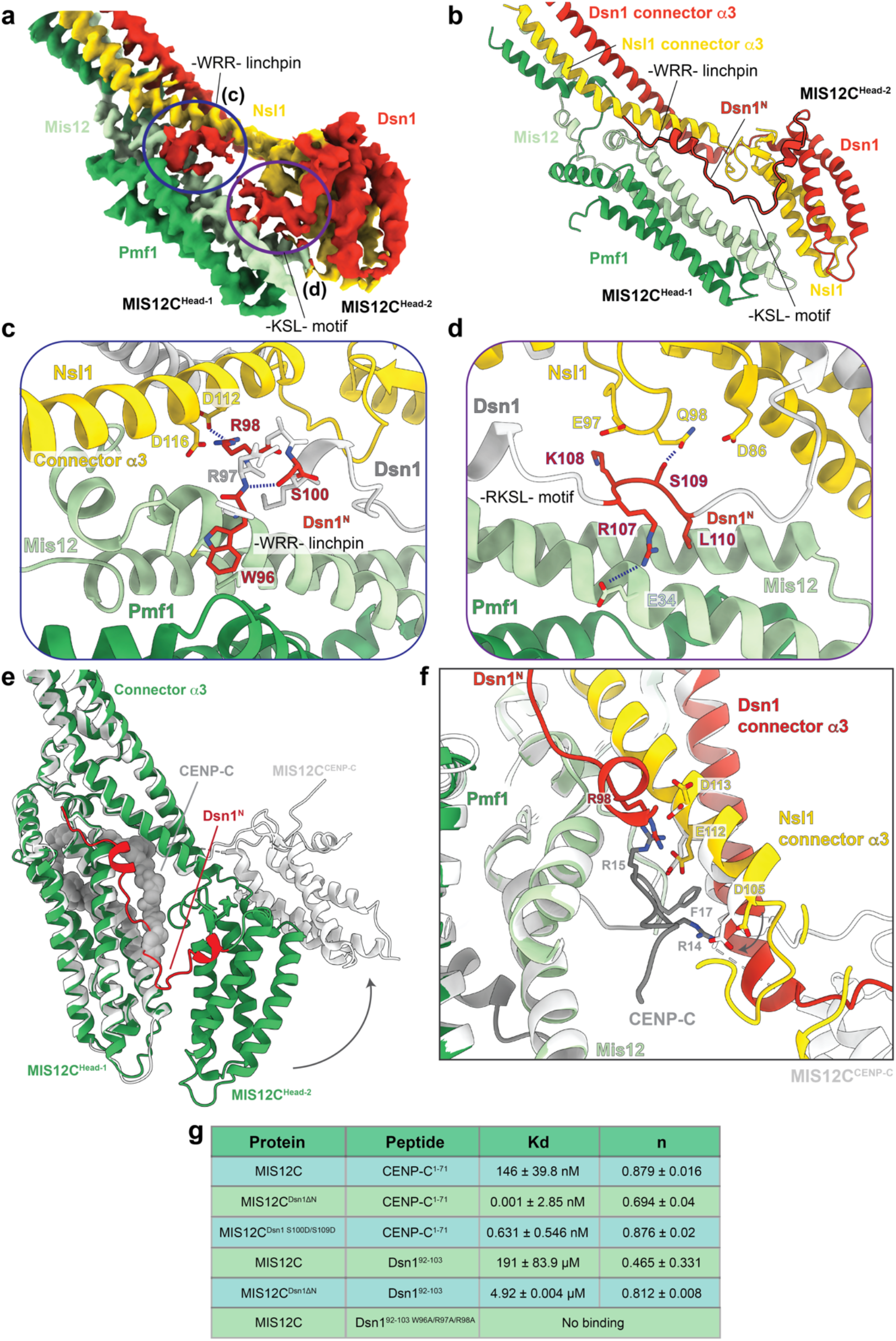
Molecular details of MIS12C complex auto-inhibition. a. Cryo-EM density map of MIS12C^Head-1^ and MIS12C^Head-2^ in the auto-inhibited state. b. Molecular model of the MIS12C head groups in the auto-inhibited state, highlighting the auto-inhibitory Dsn1^N^ element. c. Molecular details of the -WRR-linchpin interaction with the MIS12^Head-1^, with Ser100 forming stabilizing interactions with the backbone of the linchpin loop. d. Molecular details of the -RKSL-zipper binding across MIS12C^Head-1^ and MIS12C^Head-2^ and locking the auto-inhibited MIS12C state. e. The MIS12C:CENP-C complex structure (grey, PDB ID: 5LSK^11^ overlayed over the auto-inhibited MIS12C structure (green, this work). CENP-C is displayed in space-filling representation and Dsn1^N^ is coloured in red, showing a direct steric clash between CENP-C and Dsn1^N^. f. Rotation of MIS12^Head-2^ on transition from the MIS12 auto-inhibited state to the MIS12C:CENP-C complex creates a binding site for CENP-C through removing steric hindrance from both MIS12^Head-2^ and the Dsn1/Nsl1 connector-α3 helices , which also optimizes contacts to Arg14, Arg15 and Phe17 of CENP-C. The N-termini of the Dsn1/Nsl1 connector-α3 helices bend slightly on transition to the MIS12C:CENP-C complex. g. Summary of the ITC data for dissociation constants (Kd) and stoichiometry (n) between MIS12C variants and either CENP-C^1–71^ or Dsn1^92–103^ peptides, with experimental data and detailed descriptions in Extended Data Fig. 4.

At the core of the KMN^Junction^ complex is MIS12C which comprises a tetrameric coiled-coil connected to two head-domains (MIS12C^Head-1^ and MIS12C^Head-2^) (Fig. 1a-c, Extended Data Movie 1). Compared to the previously reported MIS12C crystal structure^11^, MIS12^Head-1^ and MIS12^Head-2^ in our reconstructions are closely aligned with one another, an interaction stabilized by Dsn1^N^ that folds along the MIS12^Head-1^ surface (Fig. 1b). This generates an auto-inhibited MIS12C state with an occluded CENP-C binding site, described in detail below. Spc24:Spc25^RWD^ and Knl1^RWD^ domains are rigidly docked to the top of the MIS12C stalk, and their interactions with MIS12C are further stabilized by extensive peptidic interactions with the C-termini of the MIS12C subunits Dsn1 and Nsl1. ZWINT forms a coiled-coil together with Knl1 immediately N-terminal to the Knl1^RWD-N^ domain, consistent with this segment of Knl1 being required for ZWINT interactions *in vivo*^12^. Overall, our structure has a similar global shape to the previously reported negative stain reconstruction of the truncated KMN complex^12^, and we do not observe any direct protein-protein interactions between NDC80C and KNL1C.

A salient feature of KMN^Junction^ is the coiled-coils of Spc24:Spc25 and Knl1:ZWINT that run perfectly parallel to one another, giving the entire assembly a prong-like shape (Fig. 1b). This coiled-coil arrangement projects the microtubule-binding Ndc80:Nuf2 and N-terminus of Knl1 in the same direction and away from the MIS12C head domains that associate with the inner kinetochore. Therefore, a rigid association of MIS12C:RWD:coiled-coils results in a long, rigid structure that interacts with the inner kinetochore at one end and microtubules of the mitotic spindle at the other, defining a precise polarity between the centromere-proximal MIS12^Heads^ and microtubule-proximal Ndc80:Nuf2 coiled-coils. Although the resolution of the Spc24:Spc25 coiled-coils quickly decreases away from the central MIS12C stalk, the coiled-coils are apparent at a lower cryo-EM density threshold and in 2D class averages, allowing the Spc24:Spc25 coiled-coils to be traced up to the Ndc80:Nuf2 tetramerization junction (Fig. 1d, e).

## Dsn1 N-terminus stabilizes the MIS12C auto-inhibited state

In our reconstruction, MIS12^Head-2^ directly contacts MIS12^Head-1^, resulting in an auto-inhibited MIS12C state (Fig. 2a, b; Extended Data Fig. 3a). This state is stabilized by Dsn1^N^ which forms two contacts with MIS12^Head-1^: (i) a linchpin formed by the -WRR-motif (residues 96-98) of Dsn1^N^ that locks into the hydrophobic pocket at the Mis12:Pmf1 interface and connects Mis12:Pmf1 to the coiled-coil connector-α3 helices of Nsl1:Dsn1 (Fig. 2a-c); (ii) a zipper-like interaction of the -RKSL-motif (residues 107-110) that bridges MIS12^Head-1^ and MIS12^Head-2^ through both electrostatic and non-polar interactions (Fig. 2a, b and d; Extended Data Fig. 3a). The -WRR-linchpin is additionally stabilized by a short α-helix immediately to its C-terminus, with Ser100 forming interactions with the peptide backbone of the linchpin and holding it in place (Fig. 2c). Lastly, we also observe a direct interaction between the tips of Nsl1 and Mis12 across the two heads, additionally stabilizing the auto-inhibited state (Fig. 2c).

**Figure 3.**
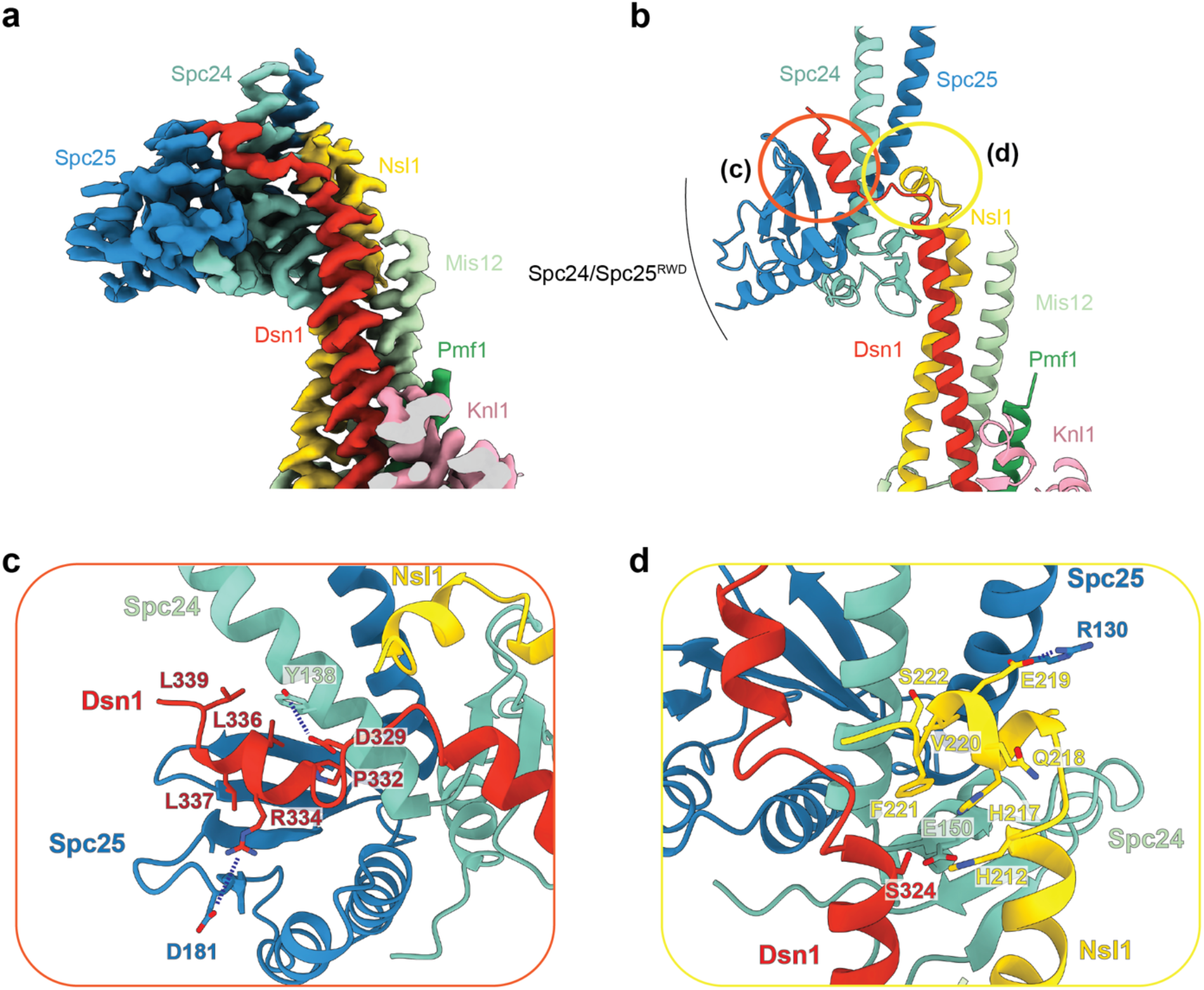
Spc24:Spc25 forms a rigid interface with MIS12C. a. Cryo-EM density map of the MIS12C:Spc24:Spc25 interface. b. Molecular model of the MIS12C:Spc24:Spc25 interface, composed of rigidly docked Spc24:Spc25^RWD^ formed from both a Dsn1 peptidic contact with Spc24:Spc25 and Nsl1 peptidic interface with Spc25. c. Molecular details of the Dsn1 peptidic interaction with Spc24:Spc25, highlighting a number of electrostatic interactions and a buried hydrophobic surface of Dsn1. d. Molecular details of the Nsl1 peptide interaction with Spc24:Spc25, similarly showing an electrostatic and geometric match between the Nsl1 peptide and Spc25 surface.

The cumulative outcome of these interactions is that the two MIS12C head domains are closely associated with little accessible space between them. This effectively blocks access to the N-terminal half of the CENP-C binding interface on MIS12^Head-1^ (Fig. 2e). Overlaying the MIS12C:CENP-C crystal structure^11^ onto our auto-inhibited MIS12C structure, we observe a direct clash between CENP-C and Dsn1^N^ in addition to the steric clash of MIS12^Head-2^ and the Dsn1/Nsl1 connector-α3 helices and parts of CENP-C (Fig. 2e). The N-termini of the Dsn1/Nsl1 connector-α3 helices undergo a modest bend on transition from the MIS12C auto-inhibited state to the MIS12C:CENP-C complex that contributes to creation of the CENP-C-binding site (Fig. 2e, f). Overall, in the MIS12C auto-inhibited state, CENP-C residues 6-22 are precluded from binding to MIS12^Head-1^, while the CENP-C residues 28-48 can still interact with the back of the MIS12^Head-1^ (Extended Data Fig. 3b, c). Thus, while MIS12C auto-inhibition would not completely abolish an interaction between CENP-C and MIS12C, it will significantly reduce the binding affinity between these two partners. Complete CENP-C binding would require disengagement of Dsn1^N^ and opening of the MIS12C central channel by repositioning MIS12^Head-2^ outwards, as observed in the MIS12C:CENP-C crystal structure^11^. This relieves steric clashes between CENP-C residues His8 to Asn11 and Arg14 to Phe17 with MIS12^Head-2^, and concomitantly positions Asp105, Glu112 and Asp113 of the Nsl1 connector-α3 helix to form hydrogen bonds with Arg14 and Arg15 of CENP-C (Fig. 2f). In the MIS12C:CENP-C complex^11^, the guanidinium group of CENP-C Arg15 occupies a similar position to that of Arg98 of Dsn1^N^ in the MIS12 auto-inhibited structure (Fig. 2f). Additionally, the shifted Dsn1/Nsl1 connector-α3 helices optimally position a non-polar pocket to engage the aromatic side chain of CENP-C Phe17 (Fig. 2f). Strikingly, auto-inhibition via Dsn1^N^ is stabilized by two key targets of the Aurora B kinase^17, 33, 36, 37^: Ser100 stabilizes the -WRR-linchpin (Fig. 2c), while Ser109 stabilizes the -RKSL-zipper (Fig. 2d). Our modelling suggests that phosphorylation of either of these residues would significantly destabilize Dsn1^N^ and would alleviate MIS12C auto-inhibition.

We used isothermal titration calorimetry (ITC) to test this structural model by measuring the affinities between MIS12C and either CENP-C or the Dsn1^N^ inhibitory peptide, including structure-guided interface mutants (Fig. 2g; Extended Data Fig. 4). We observed that the N-terminal region of CENP-C (residues 1-71; CENP-C^1–71^) bound to MIS12C with an affinity of ∼150 nM, a value similar to previous fluorescence polarization-based measurements^11, 14^. Deletion of the auto-inhibitory Dsn1^N^ segment from MIS12C significantly enhanced the MIS12C interaction with CENP-C^1–71^, with the affinity estimated to be in the sub-picomolar range, indicating an extremely tight interaction. Introducing phospho-mimetic mutations at the key regulatory Aurora B phosphorylation sites of Dsn1^N^ (S100D and S109D, MIS12^Dsn^^1^ ^S100D/S109D^) resulted in a 230-fold increase in MIS12 affinity for CENP-C^1–71^ (Kd of 0.6 nM; Fig 2g; Extended Data Fig. 4c). Our data agree with the observation that deleting a ten-residue segment of Dsn1 incorporating Ser100 and Ser109 increased by 60-fold the affinity of MIS12C for residues 1-21 of CENP-C^11^, although a smaller increase in CENP-C binding to a phospho-mimetic mutant of MIS12 was also reported^36^. Consistent with these *in vitro* results, mutating Ser100 and Ser109 to Ala, which prevents their phosphorylation, significantly reduces Dsn1 localization to kinetochores in human cells, while introducing equivalent mutations in yeast results in loss of viability^36–38^, suggesting that auto-inhibition is a physiologically relevant and important step in licensing outer kinetochore assembly. We note that another study observed a more modest effect on Dsn1 localization in human cells with Ser100 and Ser109 mutated to Ala, when combined with additional Ser to Ala mutations at the extreme Dsn1 N-terminus that we cannot model in our structure^33^. Lastly, we tested our model of the Dsn1^N^ auto-inhibitory peptide interactions with MIS12C heads. Consistent with our structure and model, the auto-inhibitory peptide of Dsn1 (residues 93-103; Dsn1^93–103^) bound to MIS12C with micromolar affinity, an interaction that was significantly enhanced by deleting the auto-inhibitory Dsn1^N^ element from MIS12C (MIS12C^Dsn^^1^^ΛN^) (Fig. 2g; Extended Data Fig. 4d, e). The interaction of Dsn1^93–103^ with MIS12C was completely abolished by mutating the -WRR-linchpin of the Dsn1^93–103^ peptide to Ala, demonstrating that the -WRR-motif is essential for auto-inhibitory Dsn1^N^ peptide binding (Fig. 2g; Extended Data Fig. 4f).

**Figure 4.**
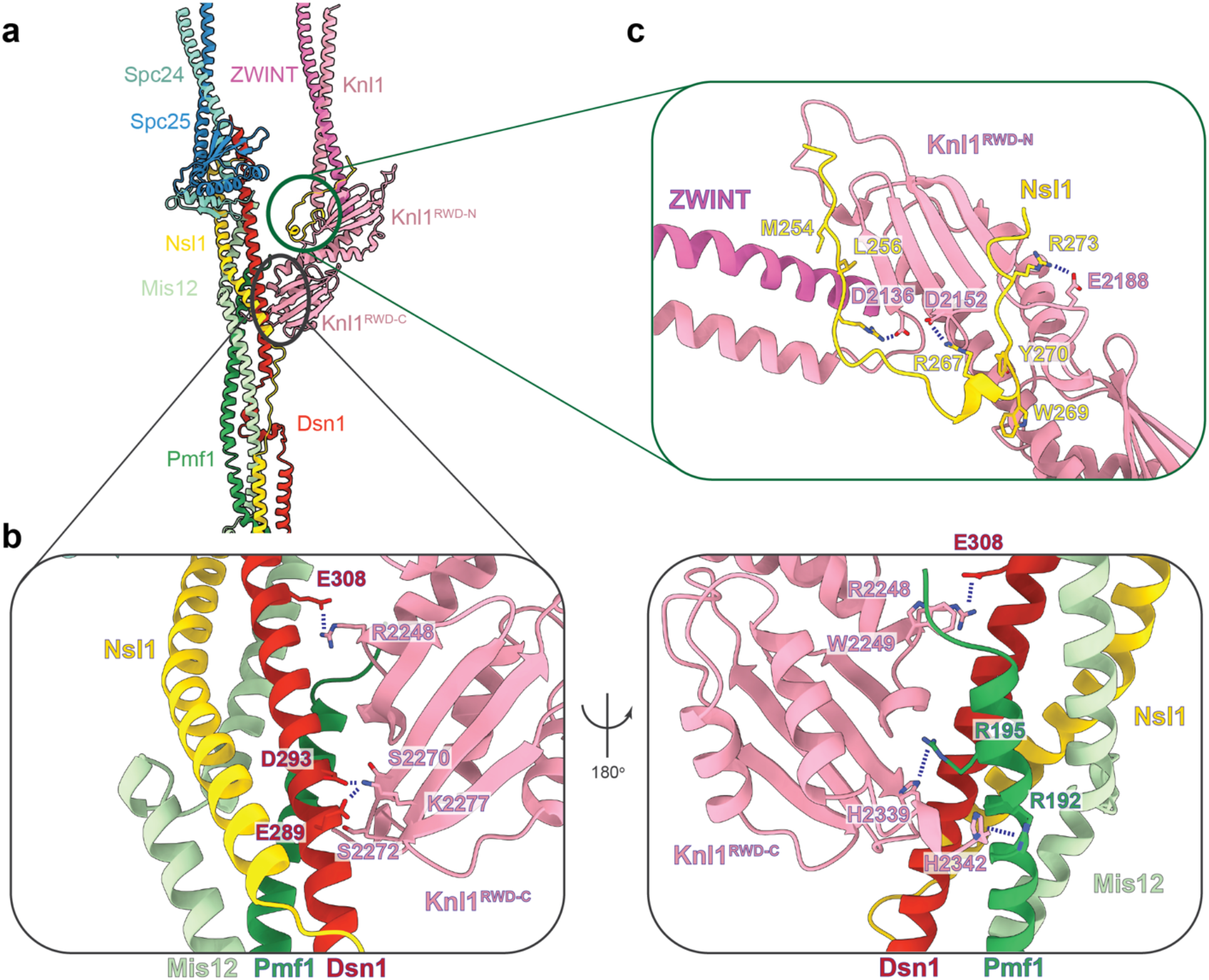
KNL1C forms multiple interactions with MIS12C. a. Overview of the MIS12C:KNL1C interface. The pair of RWD domains of Knl1 dominate this interface. b. Molecular details of the Knl1^RWD-C^ interaction with MIS12C stalk, formed by Trp2249 docking into shallow grooves of the MIS12C central stalk and additionally supported by a number of electrostatic interactions on the opposite end of the Knl1^RWD-C^. c. The Nsl1 C-terminal peptide augments the central Knl1^RWD-C^ β-sheet interaction with MIS12C, forming additional contacts with Knl1^RWD-N^.

To further assess the role of the auto-inhibitory Dsn1^N^ on CENP-C binding to MIS12C in solution, we used analytical size-exclusion chromatography (SEC) (Extended Data Fig. 5a, b). Wild-type MIS12C only partially bound CENP-C^1–71^, consistent with our structural model where half of the CENP-C binding interface is blocked. Relief of Dsn1 auto-inhibition to expose the complete CENP-C-binding site either by deleting Dsn1^N^ or mutating the -WRR-linchpin motif of Dsn1^N^, resulted in almost complete CENP-C binding, while incorporation of the phosphomimetic MIS12^Dsn1^ ^S100D/S109D^ mutants also strongly increased CENP-C binding. These results are in agreement with *in vitro* MISC12C:CENP-C affinity data reported here and previously^11^, localization experiments in cells^36–38^, and our structural prediction that phosphorylation of Ser100 and Ser109 relieves Dsn1^N^-mediated auto-inhibition to allow CENP-C binding.

**Figure 5.**
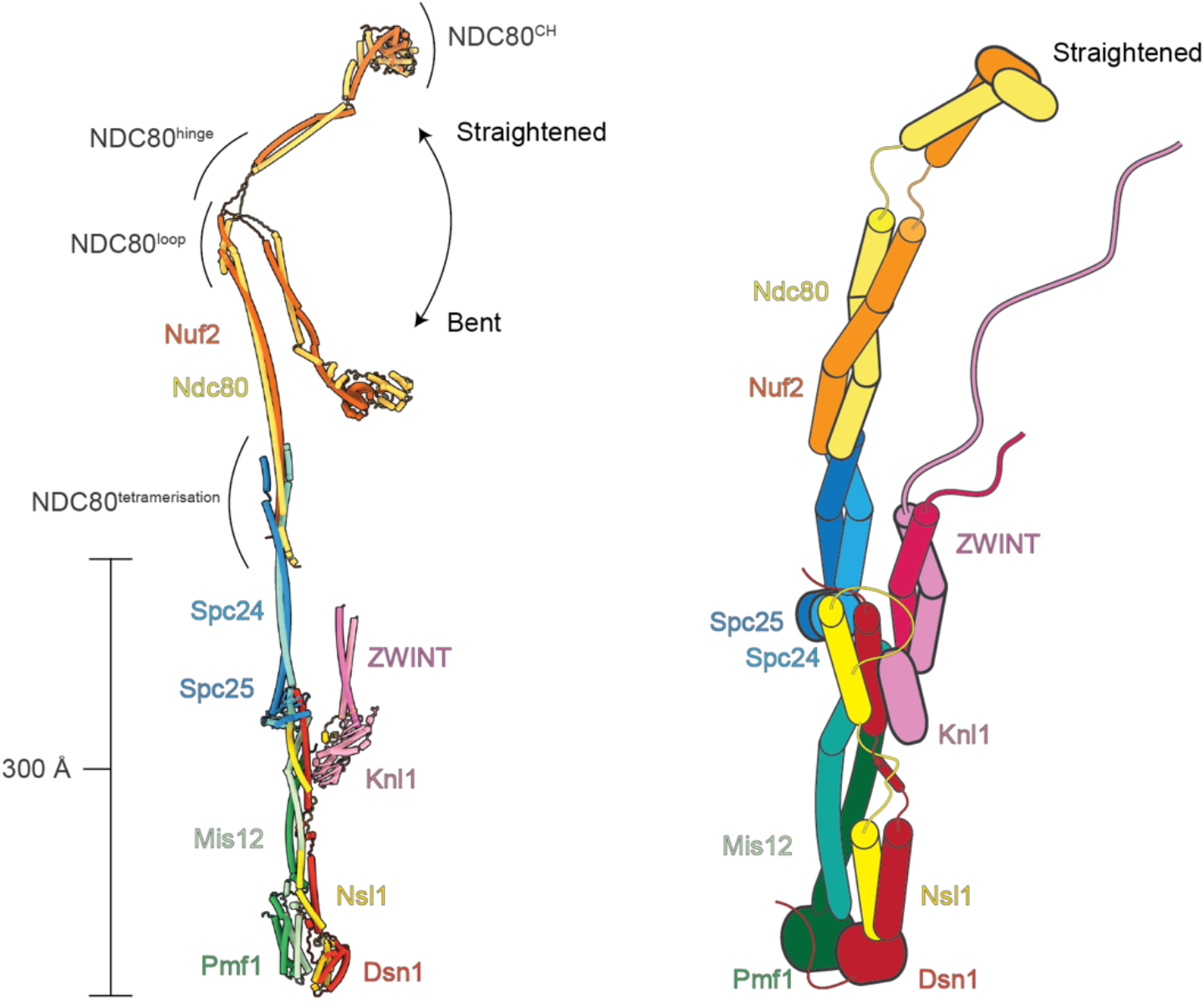
Molecular model of the complete human KMN network complex. Molecular model and cartoon schematic of the complete KMN network complex based on the structures determined in this study as well as AlphaFold2 models.

The interaction of *S. cerevisiae* MIND (the homolog of human MIS12C; termed Mis12^MIND^) with CENP-C is also regulated by Aurora B kinase phosphorylation of two equivalent serine residues in Dsn1^37, 38^. Biochemical studies of *K. lactis* Mis12^MIND^ support a model in which an unstructured region of Dsn1, incorporating the two Aurora B target sites, auto-inhibits CENP-C binding to Mis12^MIND^, and that Aurora B kinase phosphorylation of these two sites relieves this auto-inhibition^10^. To understand if the molecular basis of yeast Mis12^MIND^ auto-inhibition is evolutionary conserved with human, we used AlphaFold2^39^ to predict the structure of the full-length budding yeast Mis12^MIND^ complex. Although AlphaFold2 failed to predict the full-length *S. cerevisiae* Mis12^MIND^ complex, it could predict the related *K. lactis* Mis12^MIND^ structure (*Kl*-Mis12^MIND^) (Extended Data Fig. 5c, d). In this prediction, the *Kl*-Dsn1 N-terminus also binds across the surface of the *Kl*-Mis12^MIND^ Head-1 domain, bridging it to the Head-2 domain and occluding the hypothetical CENP-C binding interface. This predicted *K. lactis* Mis12^MIND^ complex auto-inhibition is strikingly similar to the structure we observe for human MIS12C although some of the sequence motifs involved in the interaction have diverged between the two species. This analysis suggests that MIS12C auto-inhibition is likely to be an evolutionarily conserved mechanism of regulating inner-outer kinetochore assembly in many eukaryotic lineages.

## Dsn1^N^ likely inhibits the binding of the CENP-T to MIS12C

The molecular details of the interaction between CENP-T, the other major receptor of the KMN network at the centromere, and the MIS12C complex are unknown. This limitation prevents us from unambiguously addressing, based on our structure, how MIS12C auto-inhibition also regulates CENP-T binding. Numerous lines of evidence however, would suggest that Dsn1^N^ also likely inhibits CENP-T:MIS12C association: (i) CENP-T and CENP-C cannot simultaneously bind the same MIS12C complex^18^, suggesting that they recognize an overlapping interface, and (ii) introducing mutations into MIS12C that would perturb auto-inhibition, such as deleting MIS12^Head-2^ or mutating residues in the vicinity of the -RKSL-zipper, significantly stimulated CENP-T binding^19^. In order to address this question further, we used AlphaFold2 to predict an interaction between CENP-T and MIS12C (Extended Data Fig. 6a). AlphaFold2 predicted with high confidence in all models that the CENP-T region comprising residues 185-230 specifically binds across the MIS12^Head-1^ domain and parts of the MIS12C coiled-coils (Extended Data Fig. 6a), consistent with observations that this region is essential for CENP-T-based MIS12C recruitment to kinetochores in cells^15, 17, 40^. A peptide based on this region also interacts with MIS12C *in vitro*^19^. Additionally, in the AlphaFold2 model, two key targets of CDK phosphorylation on CENP-T, Thr195 and Ser201, are in close proximity to basic residues of MIS12C, explaining how their phosphorylation would promote MIS12C:CENP-T interaction^15, 17^ (Extended Data Fig. 6b). The CENP-T binding site on MIS12C overlaps with Dsn1^N^ (Extended Data Fig. 6c), demonstrating that MIS12C auto-inhibition indeed also likely inhibits CENP-T binding, as suggested from prior *in vitro* data^19^. Notably, Arg209 of CENP-T occupies the same position as Arg98 of Dsn1^N^ (part of the -WRR-linchpin) as well as Arg15 of CENP-C, indicating that all three peptidic regions recognise a similar small site on MIS12C. Overall, MIS12C auto-inhibition also likely regulates CENP-T-based centromere recruitment of MIS12C through a comparable mechanism to that of CENP-C.

## Spc24:Spc25 forms a composite interface with MIS12C

Multiple motifs of MIS12C are important for mediating its interaction with NDC80C, particularly the C-termini of Dsn1 and Nsl1^11, 12^, with a similar Dsn1:NDC80c interaction conserved in yeast^10, 29^. Consistent with these results, we observe an extensive interaction interface between the Spc24:Spc25^RWD^ domains with the C-terminal coiled-coils as well as peptidic regions of Dsn1 and Nsl1 (Fig. 3a, b). Specifically, Spc24^RWD^ rigidly docks onto the top of the Dsn1:Nsl1 coiled-coil, which is additionally stabilized by the Mis12 α-helix on the back side (Fig. 3b). The C-terminal region of Dsn1 folds across Spc25^RWD^ (Fig. 3c), while Nsl1 forms an α-helix abutting the Spc24:Spc25 coiled-coils (Fig. 3b, d). Overall, Dsn1:Nsl1 forms an extended and intertwined connection with Spc24:Spc25, secured with multiple points of contact which might be necessary to withstand forces of the mitotic spindle across the MIS12C:NDC80C interface. Consistent with our structural model, mutating either the Dsn1 contact site (Dsn1^P332W/R334A/L336R^ mutant) (Fig. 3c) or the Nsl1 contact site (Nsl1^E219R/V220R/F221A^ mutant) (Fig. 3d) reduced the affinity of MIS12C for NDC80C as assessed by SEC experiments (Extended Data Fig. 7a, b). Combining the Dsn1^P332W/R334A/L336R^ and Nsl1^E219R/V220R/F221A^ mutants to disrupt both interfaces completely abolished MIS12C – NDC80C interactions, consistent with two independent points of contact between MIS12C and NDC80C (Extended Data Fig. 7a, b). In the accompanying manuscript, Polley et al.^41^ show that mutating either the Dsn1 or Nsl1 interfaces that contact Spc24:Spc25 significantly reduced NDC80C localization to kinetochores.

Comparison of the available crystal structures of the CENP-T:Spc24:25^RWD^ complex to our MIS12C:Spc24:Spc25^RWD^ reconstruction shows that CENP-T binds Spc24:Spc25^RWD^ at exactly the same interface as MIS12C (Extended Data Fig. 7c), confirming that MIS12C and CENP-T would compete with one another for NDC80C binding^29, 30^. This is consistent with CENP-T and MIS12C functioning in separate pathways to recruit NDC80C independently of one another. Interestingly, chicken CENP-T effectively combines both Dsn1 and Nsl1 C-terminal regions into a single polypeptide to bind across the entire Spc24:Spc25^RWD^ (Extended Data Fig. 7c).

## Knl1^RWD^ rigidly docks onto the MIS12C

The ordered C-terminal region of Knl1 consists of a tandem pair of RWD domains that transition into the Knl1:ZWINT coiled-coil (Fig. 1b, e). Knl1^RWD-C^ rigidly docks and recognizes a shallow surface formed by the Dsn1 and Pmf1 coiled-coils while Knl1^RWD-N^ uses a canonical RWD protein-interaction groove to bind the otherwise disordered Nsl1 C-terminal tail (Fig. 4a-c, Extended Data Fig. 8a). Overall, in the context of the KMN complex, each RWD domain of Knl1 engages a distinct part of MIS12C, explaining why Knl1 in most species contains two connected RWD domains. Mechanistically, this would provide robustness to the Knl1:MIS12C interaction: if one of the connections fails, the Knl1:MIS12C complex will likely remain intact because of the second redundant and independent linkage.

Knl1^RWD-C^ complements the surface and charge of the MIS12C tetrameric coiled-coil (Fig. 4a, b). Specifically, Arg2248 and Trp2249 of Knl1 insert into a pocket formed by the Dsn1 and Pmf1 subunits, an interaction that is additionally stabilized by Dsn1^E308^ and C-terminal tail of Pmf1 (Fig. 4b). On the opposite end, Knl1^RWD-C^ uses a series of electrostatic locks to engage the complementary surface charges of MIS12C (Fig. 4b).

Our use of full-length proteins allowed us to observe a long peptide of the Nsl1 C-terminus bound to Knl1^RWD-N^ (Fig. 4c). In addition to the previously reported nine residue-Nsl1^266–274^ peptide bound to Knl1^RWD-N^ (Ref. ^12^), we observe that almost the entire C-terminus of Nsl1^250–277^ (28 residues) binds across the extended interface formed by Knl1^RWD-N^ and the Knl1:ZWINT coiled-coils, with part of the interactions provided directly by ZWINT. A large portion of this interaction is well resolved in our reconstruction with amino acid-side chains clearly visible (Extended Data Fig. 8a), while a portion of this interface is only seen at lower resolution in unsharpened cryo-EM maps (Extended Data Fig. 8b). To further confirm our assignment and modelling, we used AlphaFold2 to predict the Knl1:ZWINT:Nsl1^C-terminus^ complex (Extended Data Fig. 8c). In five out of five models, the AlphaFold2 model matched our experimental structure with high confidence, including the ZWINT coiled-coil modelling, further validating our assignment. Overall, Nsl1 uses two highly conserved motifs at its C-terminus to bind to the Knl1:ZWINT complex: a -YPLR-motif binds to Knl1^RWD-N^, while a -VLKRK-motif binds across the Knl1:ZWINT coiled-coil (Extended Data Fig. 8a, b and d). The Nsl1 C-terminus buttresses the Knl1:ZWINT coiled-coils, likely stabilizing the rigid projection of these coiled-coils away from the junction.

Deletion of the Nsl1 C-terminus (MIS12C^Nsl1ΔC^) that engages Knl1^RWD-N^ significantly reduced the interaction between Knl1 and MIS12C as assessed by SEC assays, whereas in contrast, mutating the Knl1^RWD-C^ surface (Knl1^mut1^: R2248D/W2249A and Knl1^mut2^: S2270R/S2272W) had a more modest effect on reducing the affinity between these two complexes (Extended Data Fig. 9). This observation agrees with prior data that the affinity of Knl1 for the whole MIS12 complex is only slightly lower than its affinity for a peptide modelled on the C-terminus of Nsl1 alone (residues 258-281), indicating that Knl1^RWD-N^ dominates Knl1:MIS12C interactions^12^. We found that interfering with both interfaces between MIS12C and Knl1 simultaneously completely abolished the interaction between MIS12C and Knl1 (Extended Data Fig. 9), consistent with our structural model revealing two points of contact between MIS12C and KNL1C.

Previous work showed that disrupting the Knl1^RWD-N^ interaction with the Nsl1 C-terminus in cells abolished localization of Knl1 to kinetochores^12^. The accompanying study^41^ shows that mutating the Knl1^RWD-C^ surface to prevent Knl1 interactions with the MIS12C stalk also prevents Knl1 localization to kinetochores. Together, these results indicate that both Knl1^RWD-N^ and Knl1^RWD-C^ are necessary for Knl1 recruitment to kinetochores in cells.

## Overall model of the KMN network

In our cryo-EM reconstructions, we could observe only the well-resolved and conformationally rigid KMN^Junction^ module that includes the C-terminal RWD domains and coiled-coil of Spc24:Spc25 (Fig. 1a-c). The Spc24:Spc25-Ndc80:Nuf2 tetramerization junction together with the Ndc80:Nuf2 subunits were not visible in cryo-EM density (Fig. 1d, e). This could be due to either conformational heterogeneity or denaturation of these regions on the cryo-EM grid. Moderate flexing of the Spc24:Spc25 coiled-coil combined with minor conformational variability at the Spc25:Spc25^RWD^-MIS12C interface could account for the gradual reduction of cryo-EM density signal towards the Spc24:Spc25-Ndc80:Nuf2 junction, some 150 Å from the Spc24:Spc25^RWD^-MIS12C interface. Conformationally heterogeneity would be consistent with rotary shadowing EM of NDC80C, which showed both long, straight particles as well as bent particles^18^, and also consistent with the measurement of the Ndc80:Nuf2 bending in cells and in solution^34, 42^. In order to model a complete KMN network complex, we used previously published structures^22, 24, 25^, an AlphaFold2 prediction of Ndc80:Nuf2, and the structure of the KMN^Junction^ determined at high resolution in this work (Fig. 5). AlphaFold2 predicted a bent state for the Ndc80:Nuf2 complex, with the hinge region mediating bending. We could extrapolate this model to the fully straightened state, allowing us to arrive at two complete models of the human KMN network complex with the majority of details known at high resolution (Fig. 5). Our model is remarkably consistent with outer kinetochore dimensions measured in cells^34, 43^. The NDC80C bent conformation matches the KMN dimensions measured in cells in the absence of microtubule tension whereas the extended NDC80C conformation aligns well with the microtubule-bound KMN state. In our model, KMN^Junction^ represents a rigid, 300 Å long central scaffold upon which more flexibly tethered components assemble. We speculate that having a central rigid scaffold with defined dimensions is important to introduce a set distance between the chromatin proximal MIS12C complex and microtubule-proximal NDC80C in order to ensure precise positioning of the error correction and SAC components with respect to their regulation targets. For example, phosphorylation of targets of the major error correction kinase Aurora B located at the inner kinetochore is exquisitely dependent on the distance from the kinase^44^. KMN^Junction^ would provide a defined separation of Aurora B targets away from the kinase, enabling a robust error-correction function by spatial separation.

## Discussion

We determined the cryo-EM structure of the central component of the human outer kinetochore which revealed a rigid prong-shaped arrangement of eight subunits of the ten subunit KMN network complex. Our results and conclusions on the architecture of the KMN network agree with the co-submitted study by Musacchio and colleagues^41^. Their study also showed that the localization of Knl1 and NDC80C to kinetochores in cells was disrupted by structure-based mutation of their respective interfaces with MIS12C. At the core of the structure is MIS12C which interacts with the microtubule-binding NDC80C and KNL1C at the tip of its tetrameric coiled-coil. These interactions rigidly project coiled-coils of Spc24:Spc25 and Knl1:ZWINT, which are perfectly parallel to one another, away from the central MIS12C stalk and inner kinetochore recruitment sites. This arrangement precisely defines the centromere-microtubule axis along the KMN network and generates defined distances and polarity between the inner and outer kinetochore components. Our structure is consistent with such distances measured between different kinetochore components in cells^34, 43^. Additionally, we observe numerous independent interactions between NDC80C and MIS12C as well as between KNL1C and MIS12C, which are likely to function redundantly and cooperatively to provide strength and robustness to the KMN network as a central load-bearing component of the outer kinetochore, explaining how it can withstand strong mitotic forces without detachment. We validated all the novel interfaces with specific point mutations using biochemical and biophysical assays, allowing us to decouple the contribution of each specific interface to KMN complex assembly. Our structural work provides the foundation for cell-biology experiments to understand the contribution of each KMN component to kinetochore and centromere function in cells.

At the centromere-proximal end of the MIS12C, we observed an auto-inhibited MIS12^Head^ domain with inaccessible inner kinetochore binding sites. The auto-inhibited state is directly stabilized by Ser100 and Ser109, two key residues phosphorylated by centromeric Aurora B^17, 33, 35–38^. Phosphorylation of both of these residues disrupts the MIS12^Head^ auto-inhibited state and allows the KMN network to bind the inner kinetochore with extremely high affinity. Therefore, our work provides a molecular explanation of how outer kinetochore recruitment is restricted specifically to centromeres with a functional Aurora B kinase. Initial weak binding between CENP-C and MIS12C allows KMN to be recruited to the inner kinetochore. However, only KMN bound to the functional centromeres would be stabilized and converted into an extremely robust linkage by removing MIS12C auto-inhibition and allowing a very tight CENP-C:MIS12 interaction to form. Thus, intrinsic MIS12C auto-inhibition allows outer kinetochore licensing specifically at the functional centromeres. We propose that this mechanism prevents formation of active kinetochores at non-centromeric loci in cells, increasing fidelity of the chromosome segregation process. Aurora B activity is also necessary for the error-correction pathway that ensures accurate chromosome bi-orientation by resetting KMN-microtubule attachments. Thus, KMN network auto-inhibition additionally ensures that complete kinetochore assembly occurs only at sites with active error-correction machinery.

We propose a structural model for the interaction of CENP-T with MIS12C, that explains how this interaction is regulated by CDK1-phosphoryation of CENP-T^15, 17^, how CENP-T and CENP-C binding to MIS12 is mutually exclusive^18^, and how Aurora B kinase phosphorylation of Dsn1^N^ relieves auto-inhibition to significantly stimulate CENP-T binding^19^. Therefore, MIS12C auto-inhibition likely evolved to regulate both CENP-C and CENP-T-based centromere recruitment pathways.

## Materials and Methods

### Cloning of the KMN network complexes

Genes encoding *H. sapiens* Ndc80, Nuf2, Spc24, Spc25, Mis12, Pmf1, Dsn1, Nsl1 were synthesized by ThermoFisherScientific with codon optimization for *Trichoplusia ni*. The coding fragments of Knl1 and ZWINT were amplified by PCR from cDNA and cloned into a pU1 plasmid^45^. NDC80C, MIS12C and KNL1C were subsequently cloned separately into a modified Multibac expression system^45^. TEV cleavable double strep II (DS) tags were added to the C-termini of Nuf2 and Pmf1. KNL1C was subsequently re-cloned to generate the Knl1 1870-2342 fragment with an N-terminal 6xHis-SNAP tag. The constructs for Dsn1^ΔN^ (6-113 residues _deleted), Dsn1_P332W/R334A/L336R_, Nsl_1-264 _, Nsl1_E219R/V220R/F221A_, Dsn1_W96A/R97A/R98A_, were cloned_ into the plasmid containing Nsl1-Dsn1 pair and the proteins were expressed in combination with the second virus encoding Mis12-Pmf1 proteins.

Synthetic genes for *H. sapiens* Knl1^2131–2337^ and Cenp-C^1–71^, fused to a C-terminal Maltose-Binding Protein (MBP) tag (with the aacgccgccagcggt linker in between), were supplied by Integrated DNA Technologies. Both genes were inserted into pET47b plasmids carrying a kanamycin selection marker, which expresses a 3C cleavable N-terminal 6xHisTag. Mutants of Knl1^2131–2337^, namely the two double mutants R2248D/W2249S and S2270R/S2272W were created with the QuikChange protocol using primers supplied by Merck.

### Expression and purification of the KMN network and CENP-C components

The baculoviruses for expression of all KMN network complexes were generated using standard protocols^45^. All the KMN network complexes were expressed individually in High-5 insect cells. The High-5 insect cell line was not tested for mycoplasma contamination and was not authenticated. Typically, 4 L of High-5 cells were infected 2.5% v/v with the P3 cell culture and the cells were harvested 48-72 h after infection by centrifugation.

The cells expressing NDC80C or MIS12C and their associated mutants were lysed with a sonicator in lysis buffer (50 mM Tris.HCl pH 8.0, 300 mM NaCl, 0.5 mM TCEP and 5% glycerol) supplemented with benzamidine, EDTA-free protease inhibitor tablets and benzonase. Clarified lysate was loaded onto a Strep-Tactin column (Qiagen), immobilized proteins washed with lysis buffer, and the complexes were eluted in a buffer containing 50 mM Tris.HCl pH 8.0, 300 mM NaCl, 1 mM TCEP, 5% glycerol with addition of 5 mM desthiobiotin (Sigma).

For the NDC80C, the Strep-Tactin eluate was diluted to a final salt concentration of 75 mM NaCl directly and loaded onto the Resource Q anion exchange column (Cytiva). The protein was eluted in 20 mM HEPES pH 8.0, 1 mM TCEP and 5% glycerol with a NaCl gradient from 75 mM to 600 mM. The peak fraction was concentrated and further purified using a Superdex 200 16/600 column (Cytiva) in gel filtration buffer (20 mM HEPES pH 7.8, 150 mM NaCl, 1 mM TCEP). The peak NDC80C fractions were concentrated and flash frozen in liquid nitrogen.

For MIS12C, the Strep-Tactin eluate was diluted with 50 mM Tris.HCl pH 7.4 to a final salt concentration of 50 mM NaCl and directly loaded onto a Resource S anion exchange column (Cytiva). The protein was eluted in 20 mM Tris.HCl pH 7.4, 1 mM MgCl2, 1 mM TCEP and 5% glycerol with an NaCl gradient from 50 mM to 500 mM. The peak fraction was concentrated and further purified using a Superdex 200 16/600 column (Cytiva) in MIS12C gel filtration buffer (20 mM HEPES pH 7.8, 150 mM NaCl, 1 mM MgCl2, 1 mM TCEP). The peak MIS12C fractions were concentrated and flash frozen in liquid nitrogen.

Insect cells expressing KNL1C were lysed with a sonicator in KNL1C lysis buffer (50 mM Tris.HCl pH 8.0, 300 mM NaCl, 0.5 mM TCEP, 10 mM imidazole, 1 mM EDTA and 10% glycerol) supplemented with benzamidine, EDTA-free protease inhibitor tablets and benzonase. Clarified lysate was loaded onto a cOmplete His-tag purification column (Roche), immobilized proteins washed with lysis buffer, and the complexes were batch eluted in a buffer containing 50 mM Tris.HCl pH 8.0, 300 mM NaCl, 0.5 mM TCEP, 10 mM imidazole, 1 mM EDTA and 10% glycerol supplemented with 300 mM imidazole (Sigma).

NP-40 was added to the Knl1-ZWINT eluate to a final concentration of 0.01% and the eluate was diluted to a final salt concentration of 150 mM NaCl and loaded immediately onto a Resource Q anion exchange column (Cytiva). The protein was eluted in 20 mM HEPES pH 8.0, 1 mM TCEP and 10% glycerol with NaCl gradient from 150 mM to 600 mM. The eluted complex was immediately flash frozen in liquid nitrogen.

CenpC^1–71^ fused to MBP tag at its C-terminus and Knl1^2131–2337^ and its mutants were expressed and purified from *E. coli*. Plasmids were transformed into *E. coli* BL21 (DE3) cells and 3x 1L cultures were induced at an OD600 of 0.6 with 0.3 mM IPTG and grown for 18 hours at 18°C. Over-expressed proteins were purified by Ni-NTA(II) affinity chromatography (HisTrap HP, Cytiva), followed by the reverse-IMAC strategy after cleavage of the N-terminal 6xHisTag using a recombinant 3C protease carrying itself a 6xHisTag. A final purification step was performed using a Hi-load Superdex 75 gel filtration column (Cytiva) equilibrated with 10 mM Tris.HCl pH 8.0 and 150 mM NaCl. The eluted proteins were immediately flash frozen in liquid nitrogen.

### Reconstitution of the KMN network complex for cryo-EM

NDC80C, MIS12C and KNL1C were rapidly thawed, mixed in equimolar amounts (1:1:1 ratio) and applied to a Superose 6 3.2/300 (Cytiva) gel filtration column in buffer containing 20 mM HEPES pH 8.0, 80 mM NaCl, 1 mM MgCl2, 1 mM TCEP. The peak fractions were analysed by SDS-PAGE and concentrated to 1 mg/mL.

### Cryo-EM grid preparation and data acquisition

3 µl of prepared complexes at a concentration of approximately 1 mg/mL was applied to UltrAuFoil 300 mesh gold R1.2/1.3 grids (Quantifoil Micro Tools), freshly glow discharged with an Edwards S150B glow discharger for 1 min 15 sec, setting 6, 30-35 mA, 1.2 kV, 0.2 mBar (0.15 Torr). The grids were then flash frozen in liquid ethane using a ThermoFisherScientific Vitrobot IV (0-0.5 s waiting time, 2 s blotting time, -7 blotting force) using Whatman filter paper 1. Cryo-EM images were collected on ThermoFisherScientific Titan Krios microscope operating at 300 keV using a Gatan K3 camera. A magnification of 81k was used, yielding pixel sizes of 1.059 Å/pixel. The images were recorded at a dose rate of 16 e^-^/px/s with 3.2 s exposure and 45 frames. The ThermoFisherScientific automated data-collection program EPU was used for data collection, with AFIS. Defocus values ranged from -1.2 to -2.4 µm, at an interval of 0.2 µm.

### Cryo-EM data processing

Micrograph movie frames were aligned with MotionCor2^46^, and contrast-transfer function (CTF) estimation was performed by CTFFIND^47^, as integrated into RELION 4.0^48^. Particle picking was performed with TOPAZ^49^. Default parameters were used for data processing unless stated otherwise.

Particles were initially subjected to 2D classification in cryoSPARC^50^ into 100 classes using 30 online EM-iterations with 400 batchsizes per class. Classes corresponding to the KMN network complex were picked and *ab initio* models were generated (using a model number of 3). *Ab initio* models were further refined individually to obtain better-defined cryo-EM densities using homogeneous refinement and standard settings. Heterogeneous refinement in cryoSPARC with the refined KMN network volume and five decoy noise volumes was performed against the entire set of automatically picked particles using default parameters. Particles corresponding to the KMN network complex were further refined using a non-uniform refinement job type. A clean particle stack was then exported back into RELION 4.0 and two successive rounds of CTF refinement and Bayesian polishing were performed, and a final, polished particle dataset was used for a final round of 3D refinement in RELION, yielding a final resolution of 3.0 Å (GS-FSC).

In order to generate a better-resolved MIS12C head group cryo-EM densities, a mask around the MIS12C head groups was generated and classification without alignment was performed in RELION 4.0 (Extended Data Fig. 2). The class with high MIS12^Head-2^ occupancy was selected and further refined, giving a reconstruction at 3.4 Å (GS-FSC). This reconstruction was further subjected to MultiBody refinement using a previously generated mask around the MIS12C head groups. This resulted in a well-resolved MIS12C head groups at a resolution of 3.8 Å (GS-FSC).

### Cryo-EM model building and refinement

AlphaFold2 models for NDC80C and KNL1C, generated using full-length sequences, were docked into the cryo-EM densities by rigid body fitting in ChimeraX^51^ and manually corrected and adjusted in COOT^52^, including rebuilding regions *ab initio*. The models were real-space refined in PHENIX^53^ against consensus and composite maps. Default parameters for structure refinements in PHENIX were used including secondary structure and Ramachandran constraints. Figures were prepared using ChimeraX^51^.

### AlphaFold2 Predictions

AlphaFold2^39^ predictions were run using versions of the program installed locally. All the sequences used are canonical proteins sequences obtained from the Uniport.

### ITC Experiments

Isothermal titration calorimetry (ITC) was performed using an Auto-iTC200 instrument (Malvern Instruments, Malvern, UK) at 20°C. Prior to the ITC runs, all components were dialysed for at least 16 h against ITC buffer: 20 mM HEPES (pH 7.5), 100 mM NaCl, 1 mM TCEP at 4°C. For each titration run, 370 µL of protein sample was used to load the calorimeter cell. Either the Nsl1^92–103^ peptide (either wild type or mutant) or CENP-C^1–71^-MBP protein were titrated into the cell consisting of one 0.5 µL injection followed by 19 injections of 2 µL each. After discarding the initial injection, the changes in the heat released were integrated over the entire titration and fitted to a single-site binding model using the MicroCal PEAQ-ITC Analysis Software 1.0.0.1258 (Malvern Instruments). Details of the concentration of components in the syringe and cell are detailed in Extended Data Fig. 4. Titrations were performed in either duplicate or triplicate.

### Assessment of mutating the MIS12C:NDC80C interfaces

To test the effect of mutating the MIS12C:NDC80C interfaces, either wild type or three mutants _of MIS12C (MIS12C_Nsl1-E219R/V220R/F221A_, MIS12C_Dsn1-P332W/R334A/L336R_, MIS12C_Nsl1-^E219R/V220R/F221A-Dsn1-P332W/R334A/L336R^) were mixed with wild type NDC80 at 4 µM each in a buffer of 20 mM HEPES (pH 7.8), 150 mM NaCl, 0.5 mM TCEP and 50 µL was loaded onto a Superose 6 size-exclusion column (Cytiva).

### Assessment of mutating the MIS12C:KNL1 interfaces

To test the effect of mutating the MIS12C:KNL1 interfaces, either wild type MIS12C or MIS12C^Nsl1ΔC^ were mixed with either wild type or two mutants of KNL1 (KNL1^R2248D/W2249A^ (KNL1^mut1^), KNL1^S2270R/S2272W^ (KNL1^mut2^)) at 4 µM each in a buffer of 20 mM HEPES (pH 7.8), 150 mM NaCl, 0.5 mM TCEP and 50 µL was loaded onto a Superose 6 size-exclusion column (Cytiva).

### Testing the effects of the Dsn1 auto-inhibitory segment on CENP-C^1–71^ binding to MIS12C

µM of CENP-C^1–71^-MBP was mixed with 4 µM of either wild type or three mutants of MIS12C _(MIS12C_ΔN-Dsn1_, MIS12_Dsn1-W96A/R97A/R98A_, MIS12C_Dsn1-S100D/S109D_) in a buffer of 20 mM HEPES_ (pH 7.8), 150 mM NaCl, 0.5 mM TCEP and 50 µL was loaded onto a Superdex S200 size-exclusion column (Cytiva).

## Extended Data Video 1

Video detailing the cryo-EM structure of the KMN^Junction^ complex, showing cryo-EM density, and the fitted models.

## Author contributions

D.B. and S.Y. designed the study and experiments. Z.Z., D.Be. and S.Y. cloned constructs. J.Y, D.Be. and S.Y. purified all proteins. S.Y. prepared the grids, collected and processed the cryo-EM data. J.Y. performed all biochemical experiments. D.B. performed all ITC experiments. D.B. and S.Y. wrote the manuscript with input from all authors.

## Acknowledgments.

We are grateful to the LMB EM Facility for help with the EM data collection, J. Grimmett and T. Darling for computing, and J. Shi for help with insect cell expression. We would also like to thank members of Barford group for their input and useful discussions, especially K. W. Muir and N. Turner for critical reading of the manuscript. We also thank Andrea Musacchio for sharing unpublished experimental observations. For the purpose of open access, the author has applied a CC BY public copyright license to any Author Accepted Manuscript version arising.

## Data and materials availability

PDB and cryo-EM maps have been deposited with RCSB and EMDB, respectively. Accession numbers are listed in Extended Data Table 1.

---

**Extended Data Figure 1.**
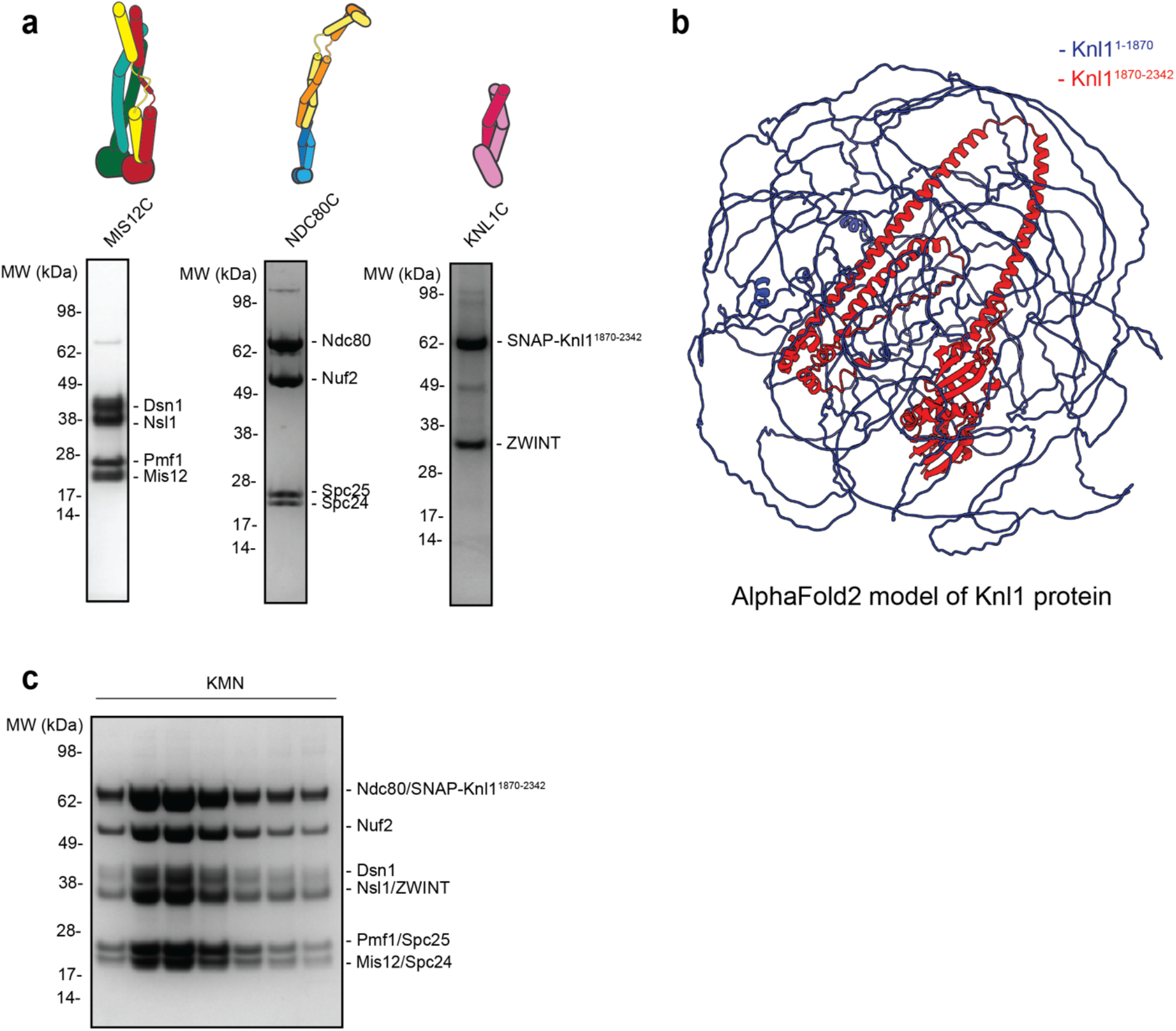
Cryo-EM sample biochemistry. a. Coomassie blue-stained SDS-PAGE gels of the purified KMN network components and their corresponding schematics that was used in this study for structure-function analysis. b. AlphaFold2 prediction of the full-length Knl1 protein obtained from Uniprot server. Knl1 region coloured in red (amino acids 1870-2342) was used in this study as it contains all structured regions of the Knl1 protein. c. Coomassie blue-stained SDS-PAGE gel of the reconstituted KMN network complex that was used for cryo-EM grid preparations and structural analysis.

**Extended Data Figure 2.**
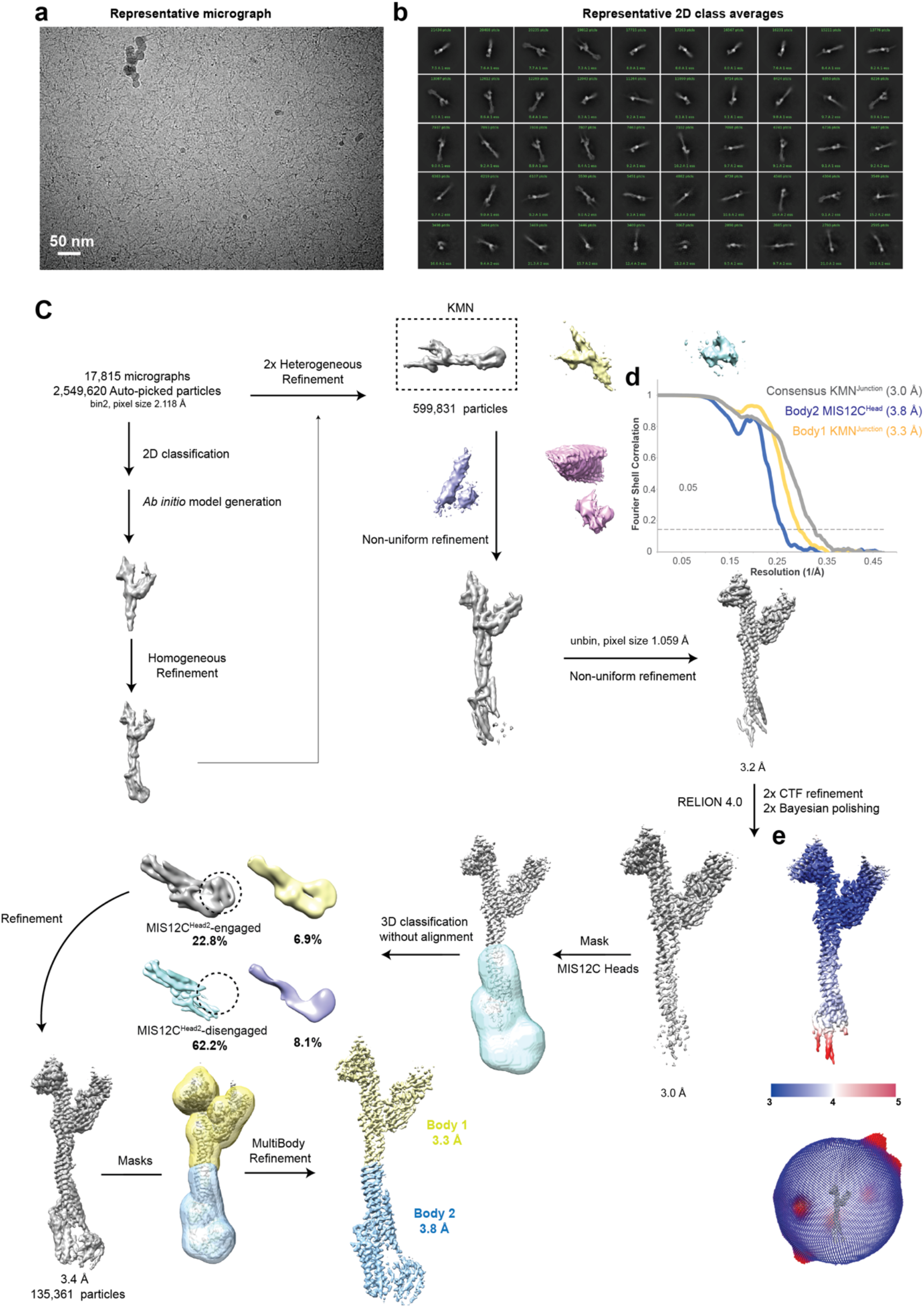
Cryo-EM data processing. a. Example of a representative cryo-EM micrograph obtained during data collection. b. Representative 2D class averages generated in cryoSPARC during initial 2D classification steps. c. Cryo-EM data processing workflow summary as described in the methods section. d. Fourier Shell Correlation (FSC) plots of the three main maps used to generate molecular models. The maps are also highlighted in (c). e. Consensus KMN^Junction^ complex coloured by local resolution as well as angular distribution of particle views.

**Extended Data Figure 3.**
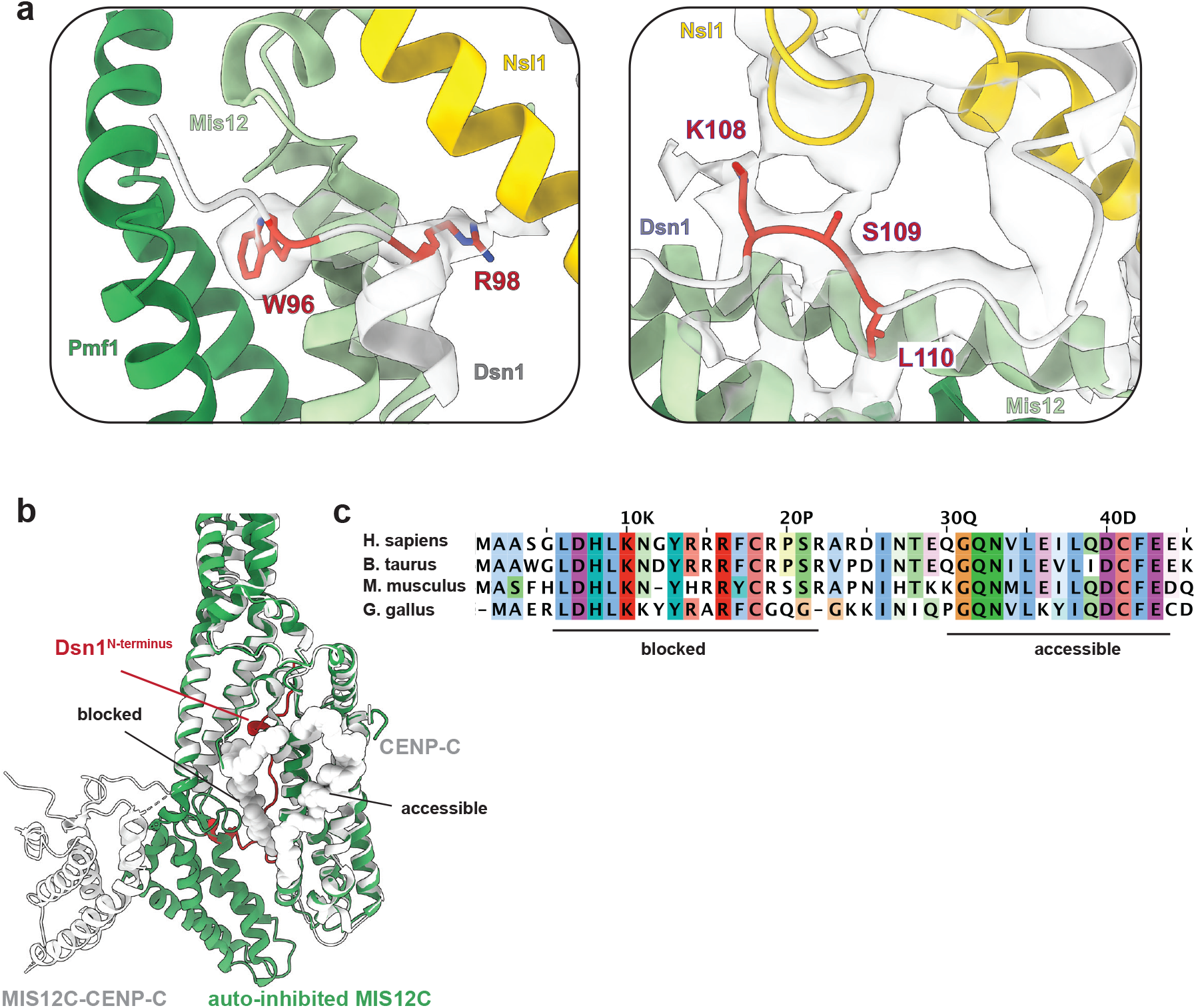
Details of Dsn1 auto-inhibition. a. Molecular model of the Dsn1^N^ fitted into the MIS12C^Head-2^ cryo-EM density map, showing side-chain resolution at the key interaction regions. b. MIS12C:CENP-C structure (grey, PDB ID: 5LSK^11^) overlayed with the auto-inhibited MIS12C complex shows that the back side of the MIS12C is available to bind a portion of the CENP-C (residues 1-21). c. CENP-C multiple sequence alignment^54^, highlighting the CENP-C regions that can bind to the MIS12C complex in the auto-inhibited state (accessible region, C-terminal portion) and the region of CENP-C (N-terminal portion) that is blocked from engaging MIS12C.

**Extended Data Figure 4.**
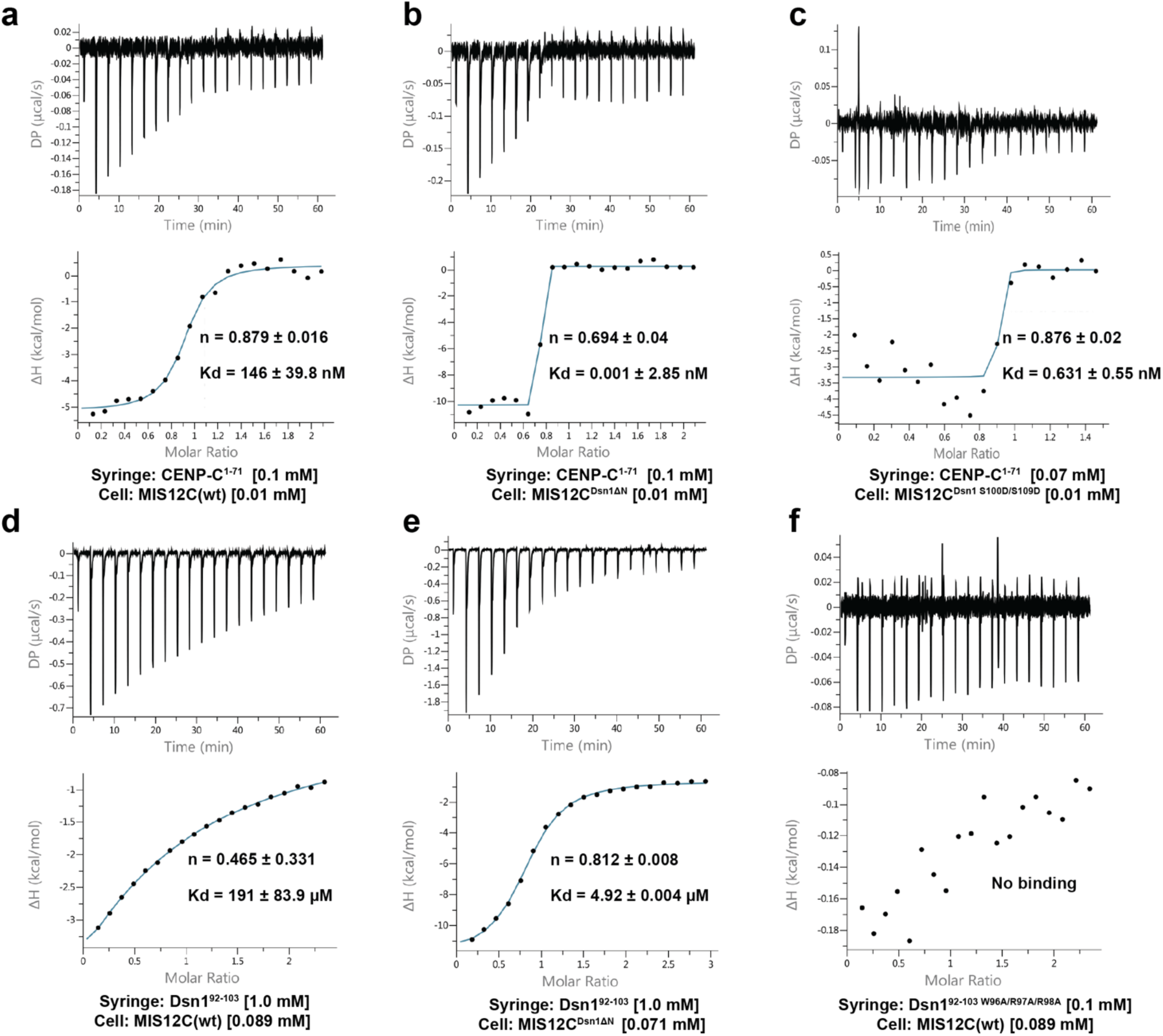
Isothermal titration calorimetry experiments. The Kd and stoichiometries (n) values are averages between at least two experiments. The reported error values are calculated standard deviations. CENP-C^1–71^ and Dsn1^92–103^ peptides were injected using a syringe into the cell containing MIS12C protein variants. Protein and peptide concentrations in cell and syringe are indicated in square brackets. a. Titration of CENP-C^1–71^ peptide to full-length and unmodified MIS12C, MIS12C (wt). b. Titration of CENP-C^1–71^ peptide to MIS12C with deleted Dsn1^N^ (6-113 residues deleted, MIS12^Dsn1ΔN^). c. Titration of CENP-C^1–71^ peptide to MIS12C with S100D and S109D substitution in Dsn1 protein (MIS12C^Dsn1^ ^S100D/S109D^). d. Titration of Dsn1^92–103^ peptide to MIS12C (wt). e. Titration of Dsn1^92–103^ peptide to MIS12^Dsn1ΔN^. f. Titration of Dsn1^92–103^ peptide with W96A/R97A/R98A substitutions (Dsn1^92–103^ W96A/R97A/R98A_) to MIS12C (wt)._

**Extended Data Figure 5.**
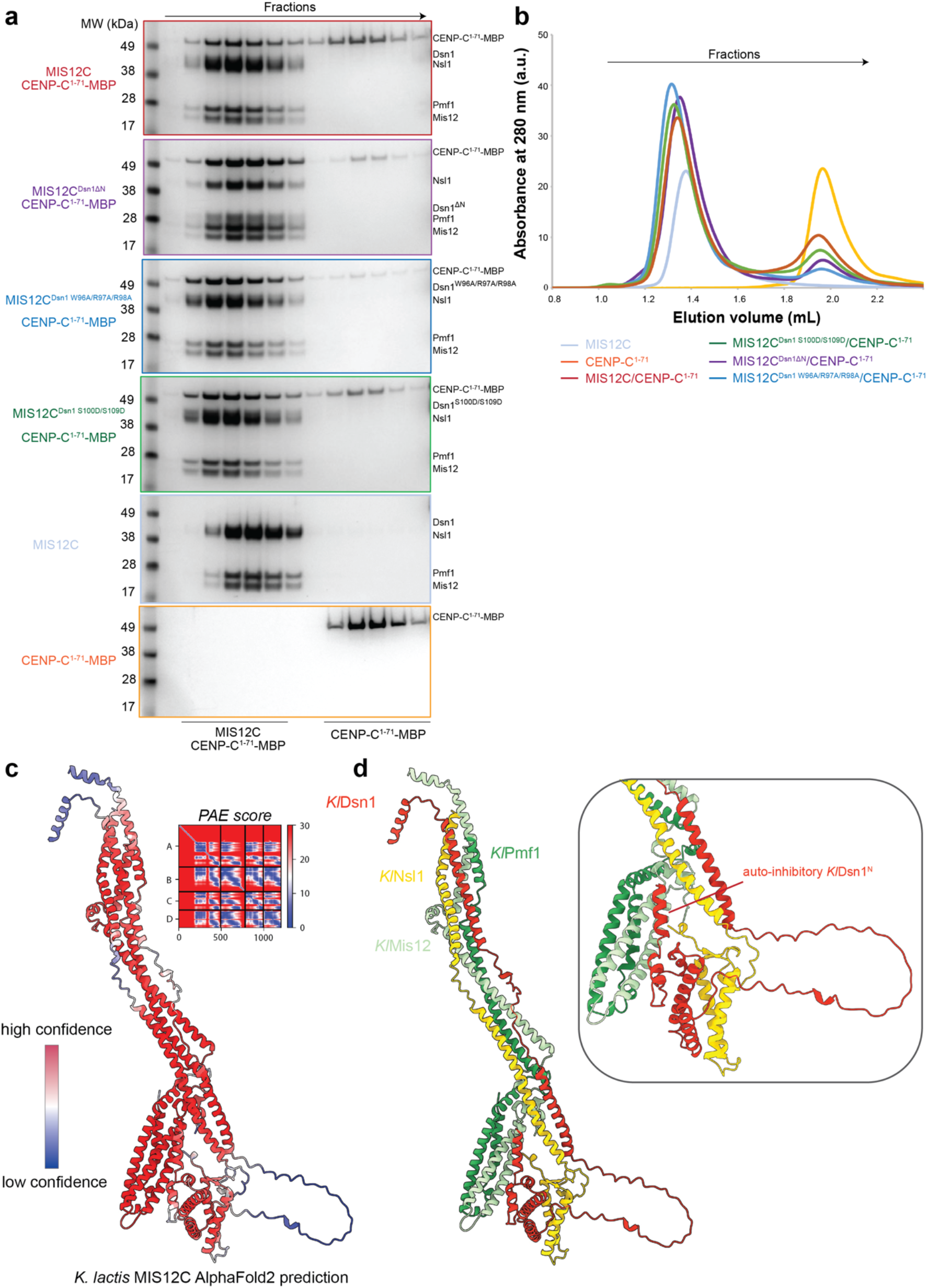
Biochemical analysis of CENP-C interactions with MIS12C. a. Coomassie blue-stained SDS-PAGE gels of the MIS12C:CENP-C^1–71^ interaction reconstitution. Wild-type MIS12C, MIS12^Dsn1ΔN^, MIS12C with W96A/R97A/R98A substitutions in Dsn1 (MIS12C^Dsn1^ ^W96A/R97A/R98A^) or MIS12C^Dsn1^ ^S100D/S109D^ was used in these experiments to test interaction with CENP-C^1–71^ tagged at the C-terminus with Maltose Binding Protein (MBP, CENP-C^1–71^-MBP) to allow robust visualization of CENP-C. b. SEC elution chromatograms of all of the experiments described in (a). c. AlphaFold2 Multimer prediction of the *K. lactis* MIS12C complex structure. Coloured by pLDDT score. d. AlphaFold2 Multimer prediction of the *K. lactis* MIS12C complex coloured by subunit with insert highlighting conservation of the auto-inhibitory Dsn1^N^ mechanism.

**Extended Data Figure 6.**
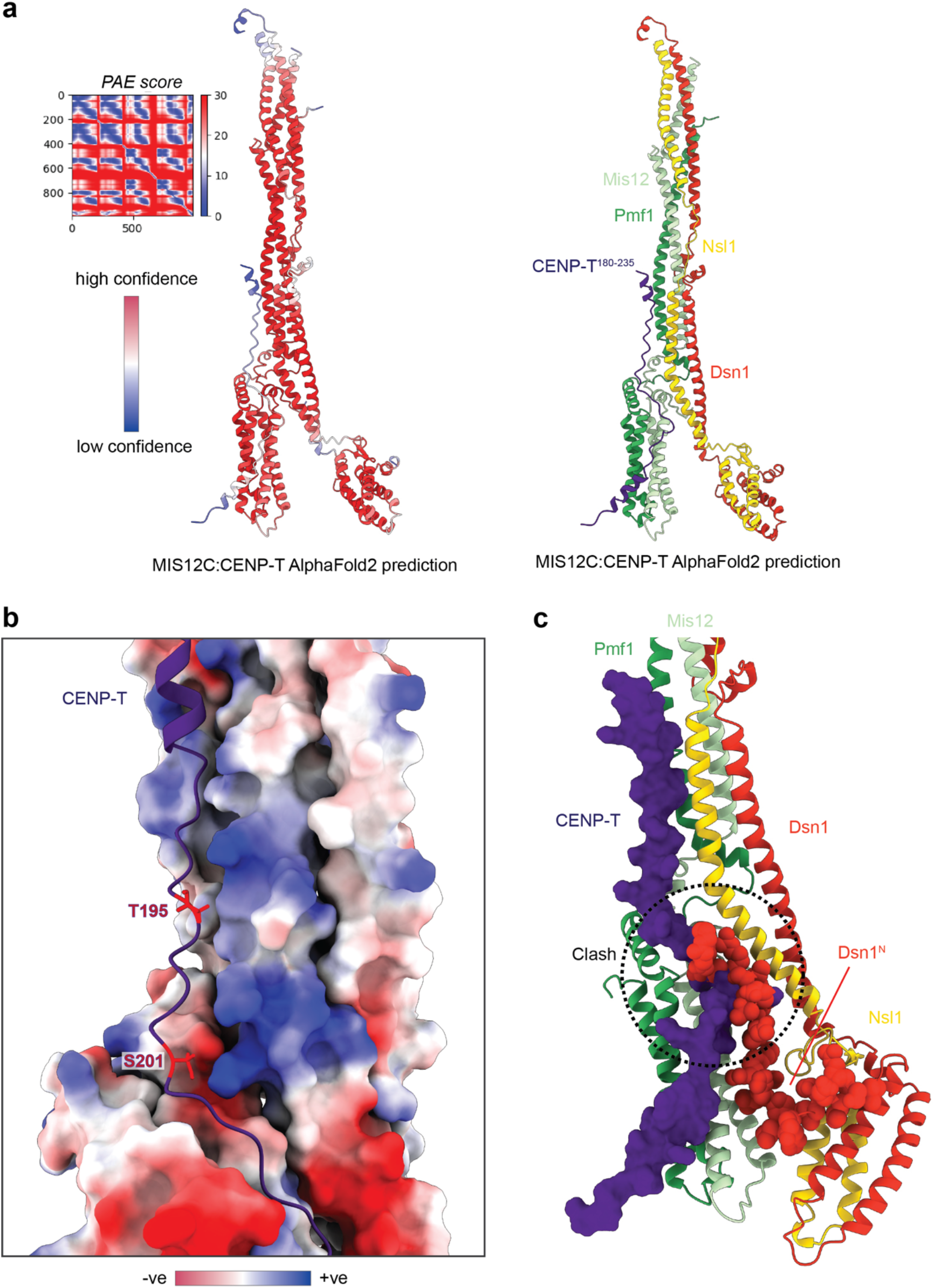
AlphaFold2 model of CENP-T:MIS12C interactions. a. AlphaFold2 Multimer prediction of the CENP-T:MIS12C interaction based on the full-length MIS12C protein sequences for all proteins apart from Dsn1, for which residues 113-356 were used to remove auto-inhibitory Dsn1^N^ region. Only CENP-T residues 180-230 were used for this prediction. b. A CENP-T-interacting region of MIS12C is shown as electrostatic surface potential while Thr195 and Ser201 of CENP-T are highlighted in red. Both Thr195 and Ser201 are targets of CDK kinase in cells and both residues face positively-charged patches on MIS12C. c. Overlay of CENP-T:MIS12C AlphaFold2 prediction on the MIS12C structure determined in this study. The auto-inhibitory Dsn1^N^ region and CENP-T peptidic region are shown in space-filling representation, highlighting an extensive steric clash.

**Extended Data Figure 7.**
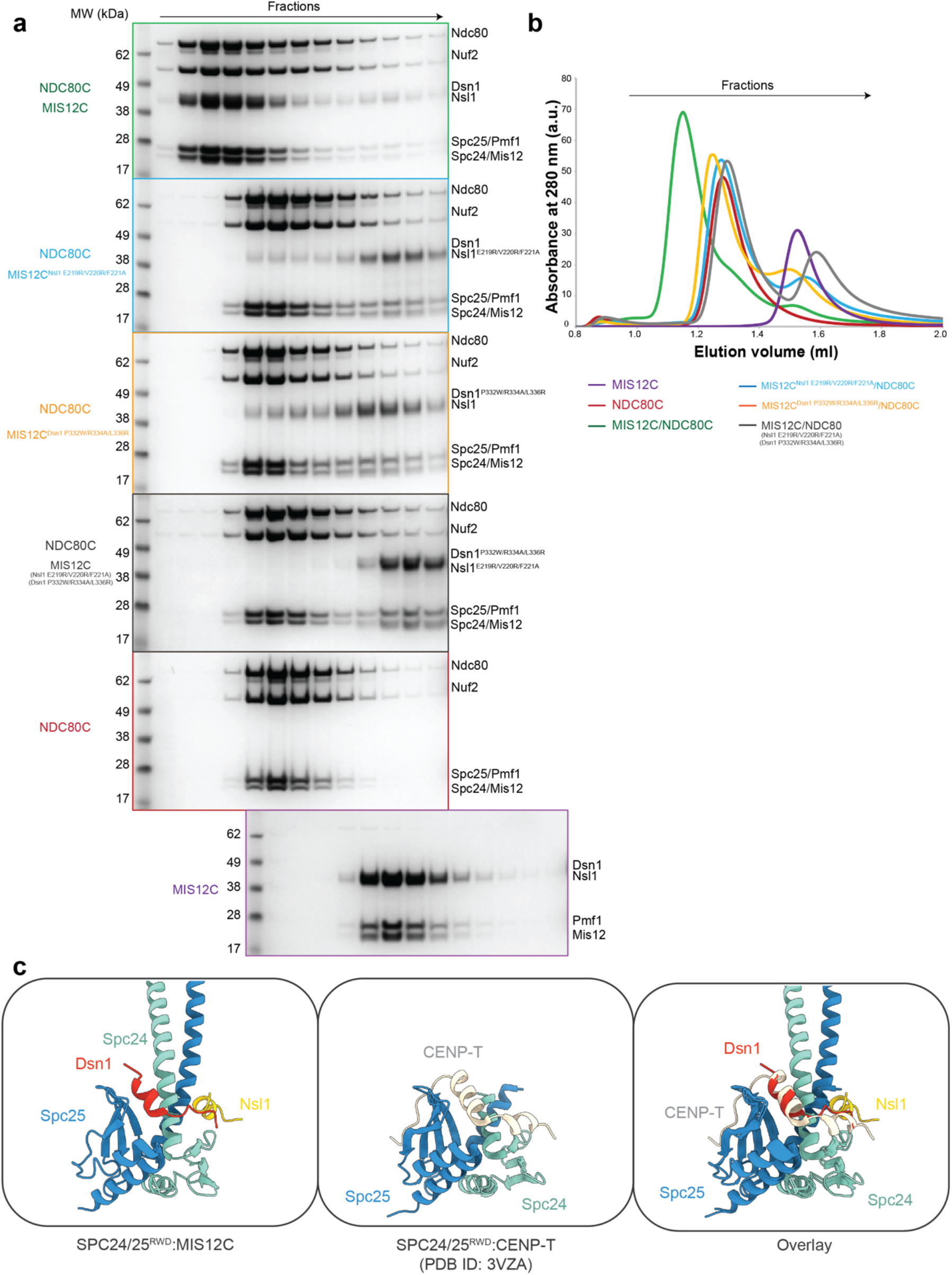
Biochemical characterization of MIS12C:NDC80C interactions. d. Coomassie blue-stained SDS-PAGE gels of the MIS12C:NDC80C interaction reconstitutions. MIS12C (wt), MIS12C with E219/RV220R/F221A substitutions in Nsl1 (MIS12C^Nsl1^ ^E219/RV220R/F221A^) and MIS12C with P332W/R334A/L336R substitutions in Dsn1 (MIS12C^Dsn1^ ^P332W/R334A/L336R^) were tested for binding to full-length, unmodified NDC80C. e. SEC elution chromatograms of all of the experiments described in (a). f. Comparison of the Spc24:Spc25 interaction with peptides from MIS12C and CENP-T (PDB ID: 3VZA^30^).

**Extended Data Figure 8.**
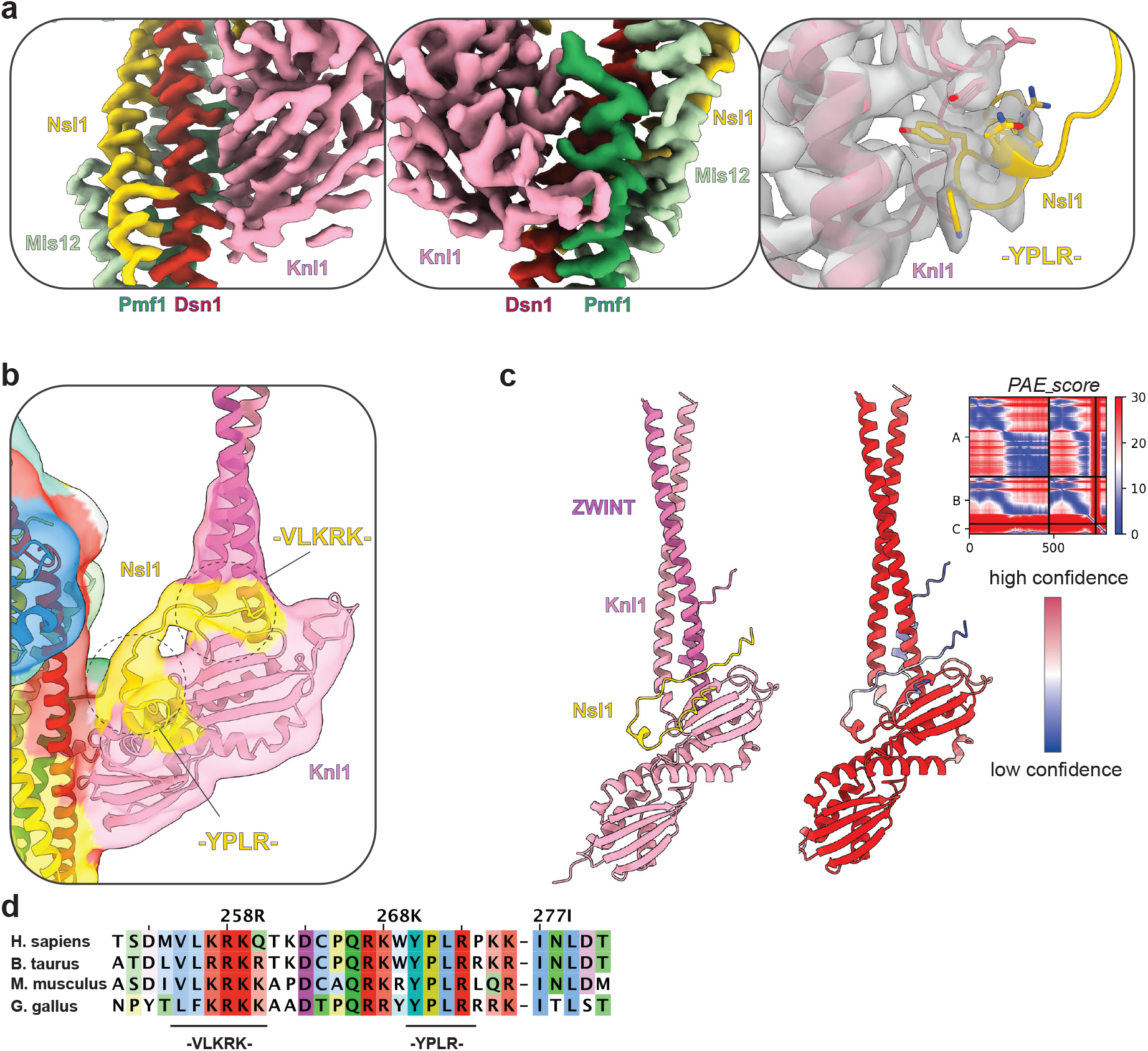
Molecular details of MIS12C:KNL1C interactions. a. Cryo-EM density map of the KNL1C:MIS12C interface, highlighting the resolution of interactions in this region. b. Unsharpened and unmasked cryo-EM map of the KMN network shows a long peptide of Nsl1 binding to the Knl1:ZWINT interface and then folding across the Knl1 surface. c. AlphaFold2 structure prediction of the Knl1:ZWINT:Nsl1 complex, demonstrating strong agreement with the experimental density maps in (b). d. Sequence alignment of the Nsl1 protein showing conservation of two Nsl1 motifs involved in Knl1 interaction.

**Extended Data Figure 9.**
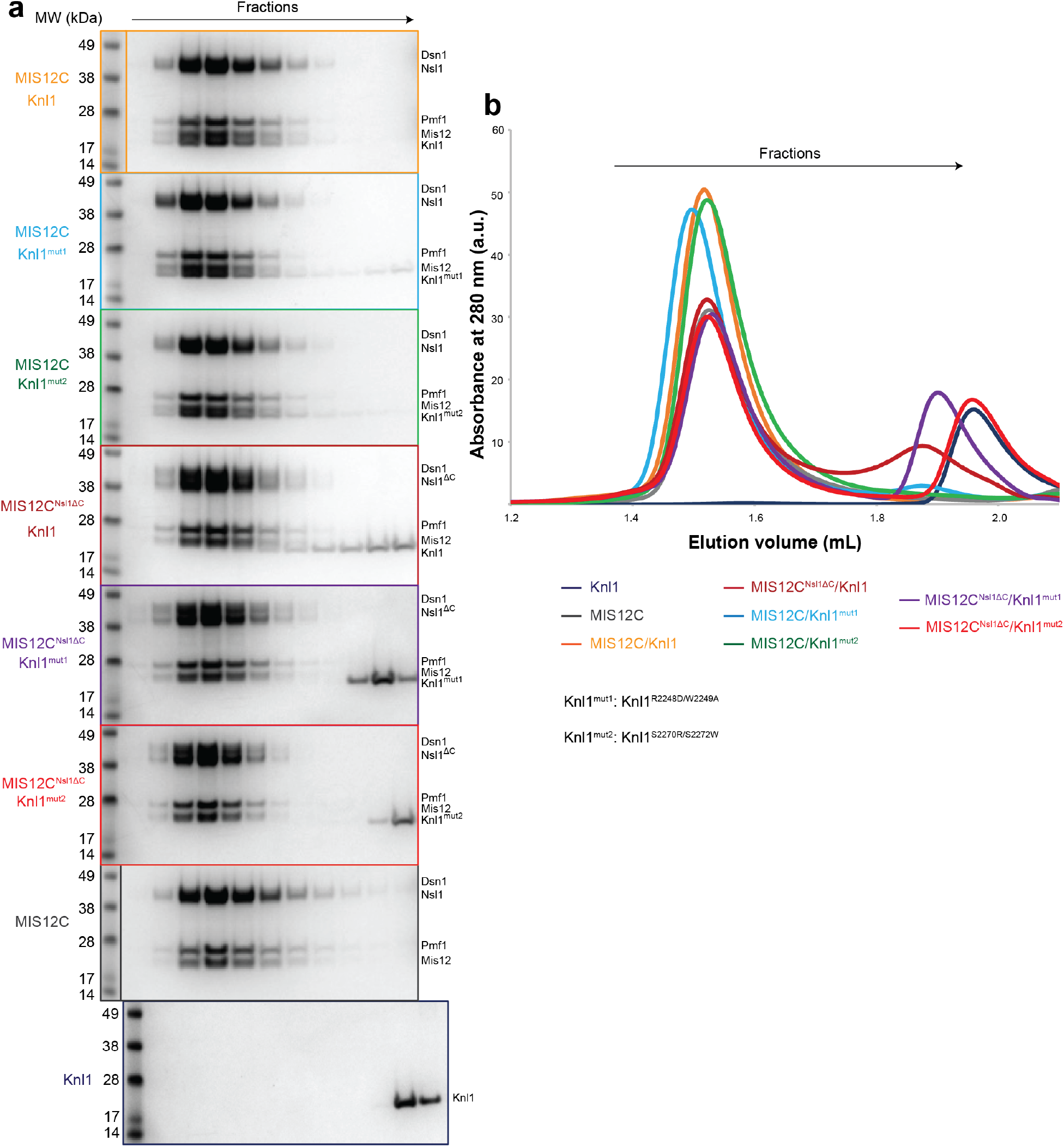
Biochemical validation of the MIS12C:Knl1 interface. a. Coomassie blue-stained SDS-PAGE gels of the MIS12C:Knl1 interaction reconstitutions. MIS12C (wt) and MIS12C with Nsl1 truncated at the C-terminus (1-264 Nsl1 construct, MIS12C^Nsl1ΔC^) were tested in their ability to bind either wild-type RWD domains of Knl1 (2131-2337 construct, labelled Knl1 for simplicity) or Knl1 mutants (R2248D/W2249S substitutions, Knl1^mut1^, and S2270R/S2272W substitutions, Knl1^mut2^). b. SEC elution chromatograms of all of the experiments described in (a).

**Extended Data Table 1.** Cryo-EM data collection, refinement and validation statistics.

|  |  |
| --- | --- |
|  | KMN network<br>complex<br>(EMDB-17814)<br>(PDB 8PPR) |
| **Data collection and processing** |  |
| Magnification | 81,000 |
| Voltage (kV) | 300 |
| Electron exposure (e<sup>-</sup>/Å<sup>2</sup>) | 45 |
| Defocus range (μm) | 1.2-2.4 |
| Pixel size (Å) | 1.059 |
| Symmetry imposed | C1 |
| Initial particle images (no.) | 2,549,620 |
| Final particle images (no.) | 599,831 |
| Map resolution (Å) | 3.0 |
| FSC threshold | 0.143 |
| Map resolution range (Å) | 2.7-20 |
| **Refinement** |  |
| Initial model used (PDB code) | *Ab initio* |
| Model resolution (Å) | 2.7 |
| FSC threshold | 0.143 |
| Model resolution range (Å) | 2.4-3.2 |
| Map sharpening *B* factor (Å<sup>2</sup>) | -61.68 |
| Model composition |  |
| Non-hydrogen atoms | 12652 |
| Protein residues | 1544 |
| Ligands | - |
| *B* factors (Å<sup>2</sup>) |  |
| Protein | 138.75 |
| Ligand | - |
| R.m.s. deviations |  |
| Bond lengths (Å) | 0.003 |
| Bond angles (°) | 0.567 |
| Validation |  |
| MolProbity score | 1.44 |
| Clashscore | 7.98 |
| Poor rotamers (%) | 0 |
| Ramachandran plot |  |
| Favoured (%) | 97.97 |
| Allowed (%) | 1.97 |
| Disallowed (%) | 0.07 |

